# The selfish yet forgetful brain: Stable cerebral oxygen metabolism during hypoglycemia but impaired memory consolidation

**DOI:** 10.1101/2024.12.12.628178

**Authors:** Antonia Bose, Stefanie J. Haschka, Johanna Koehler, Felix A. Hesse, Santiago Martin, Lea Steinberg, Roman Iakoubov, Valentin Riedl

## Abstract

The continuous supply of glucose and oxygen is essential for healthy brain function. Accordingly, the Selfish Brain Theory proposes that the human brain prioritizes its own energy demands, making it less vulnerable to fluctuations in systemic energy availability. Although studies have reported decreases in cerebral glucose metabolism, alternative energy sources other than glucose might be oxidized for ATP production. However, cerebral oxygen metabolism (CMR_O2_) has never been quantified across the human brain. In this study, we investigated the influence of insulin-induced hypoglycemia on CMR_O2_ in healthy male participants. Additionally, we explored the prolonged effects of hypoglycemia on cognitive function following the restoration of euglycemia. We found that CMR_O2_ remained stable under hypoglycemia, even at blood glucose levels below 49 mg/dL. Interestingly, we detected a significant increase in cerebral blood flow (CBF) of up to 11%, particularly in regions involved in higher cognitive processing. Despite stable rates of oxygen metabolism, we identified a selective impairment in memory consolidation following hypoglycemia, even after normal glucose levels were restored, with no effects observed in memory encoding or attention. In favor of the Selfish Brain Theory, the stability in CMR_O2_ suggests that the brain efficiently shifts to alternate energy pathways under hypoglycemia, potentially using astrocytic glycogen. Despite this metabolic flexibility, our results indicate that prior hypoglycemia imposes long-lasting effects on memory consolidation, possibly linked to glycogen depletion and impaired glutamate synthesis. In summary, our study suggests that clinical states of hypoglycemia pose a critical impact on patients’ brain health and functioning.

## Introduction

The human brain relies on oxidized glucose as its primary fuel, accounting for a disproportionally large share of the body’s total energy consumption relative to its small mass. Intriguingly, despite its high metabolic demands, the brain has minimal intrinsic energy storage and is almost exclusively dependent on the constant supply and metabolism of glucose. Unfortunately, the brain’s reaction to a reduced energy supply (e.g. glucose deficiency) remains only poorly understood. Nevertheless, better understanding of insulin-induced hypoglycemia is necessary to improve the quality of life and prognosis for insulin-treated patients (Genuth, 2006) or other patient groups in danger of hypoglycemia (e.g. insulinoma or congenital hyperinsulinism (Arnoux et al., 2010; Grant, 2005)).

Hypoglycemic blood sugar levels lead to both physical and cognitive impairments, with acute hypoglycemia often affecting attention and memory processing (Graveling et al., 2013a; McAulay et al., 2001a; Sommerfield et al., 2003a). While these effects are directly linked to glucose deficiency, insulin also plays an important regulatory role in brain metabolism. Beyond its well-documented peripheral functions, insulin signaling in the brain has been shown to control systemic energy homeostasis (García-Cáceres et al., 2016); however, the full impact of hypoglycemia on cerebral energy metabolism, as well as the specific effects of insulin, remains unknown.

The present study investigates whether cerebral energy metabolism is downregulated or remains stable under conditions of insulin-induced glucose deficiency. Initial studies in humans during hypoglycemia indicated a significant reduction of up to 30% in glucose utilization (Blazey & Raichle, 2019; Boyle et al., 1994). This suggests two potential mechanisms: (a) a general downregulation of cerebral energy metabolism, possibly reflecting an energy conservation strategy employed by the brain during fuel deficiency, or (b) a compensatory shift from oxidized glucose to the oxidation of alternate fuels to maintain metabolic homeostasis. The latter aligns with the Selfish Brain Theory, which proposes that the brain prioritizes its own energy demand (Peters et al., 2004) and would thus be resilient to fluctuations in energy substrate availability. This theory is mostly supported by the data demonstrating preserved structural and metabolic integrity of the brain during metabolic stress (Kind et al., 2005; Miller et al., 2002; Oltmanns et al., 2008; Peters et al., 2011). Unfortunately, no quantitative data on cerebral energy metabolism yet exists. Particularly, the impact of hypoglycemia on overall cerebral energy metabolism remains unclear as oxygen consumption, a more comprehensive marker of energy use, has not been directly measured under hypoglycemic conditions. Unlike glucose, which reflects the utilization of only one, albeit the primary, fuel source, oxygen consumption accounts for the oxidation of various substrates, providing a more complete picture of the brain’s metabolic activity. Our study provides novel insight into brain energy consumption and adaptation to hypoglycemic conditions by directly measuring cerebral oxygen consumption through a novel MRI-based approach.

To this end, we measured CMR_O2_ using multiparametric quantitative BOLD (mqBOLD) imaging (Christen et al., 2012; Hirsch et al., 2014; Kaczmarz et al., 2020). mqBOLD is an established MRI approach to measure absolute levels of oxygen metabolism and, therefore, offers a quantitative extension of conventional BOLD fMRI, which only captures changes in blood oxygenation. In this study, mqBOLD was combined with a hyperinsulinemic glucose clamp (Heise et al., 2016) to induce both hypoglycemia and euglycemia in healthy male humans. We only recruited male participants to control for hormonal fluctuations, particularly those associated with estrogen, which are known to influence insulin sensitivity (Yan et al., 2019). These variations could confound results when data is collected on multiple days, as was the case in our study, making it challenging to isolate the factors of interest. In addition to hyperinsulinemic hypoglycemic and euglycemic conditions, we introduced a sham condition by infusing a 0,9% sodium chloride solution, thereby allowing us to disentangle the specific effects of insulin on brain energy metabolism.

Furthermore, we examined the long-term effects of hypoglycemia on cognitive function. While cognitive impairment during hypoglycemia is well-documented (Broadley et al., 2022; Graveling et al., 2013; Sommerfield et al., 2003b), it remains unclear whether these deficits persist after euglycemic conditions are restored. We specifically investigated whether memory systems, which are critically dependent on glucose (de Tredern et al., 2021), exhibit long-term impairments following hypoglycemia, even after glucose levels return to normal.

Our data show stable rates of cerebral oxygen consumption (CMR_O2_) under hypoglycemia. This provides evidence for the notion that the brain can maintain its energy metabolism, even under conditions of glucose deficiency, consistent with the Selfish Brain Theory. Despite stable metabolic rates, our results reveal long-term behavioral effects of hypoglycemia: specifically, even when subjects learned under restored euglycemia following hypoglycemia, their performance on a memory recall task was significantly worse 24 hours later compared to continuous euglycemic conditions.

## Results

In the present study, we employed mqBOLD imaging to quantitatively assess the effects of hypoglycemia on cerebral oxygen metabolism (CMR_O2_) (Fig. 1A). To this end, we performed hyperinsulinemic glucose clamping and acquired mqBOLD MR data concurrently. Each participant underwent three clamping sessions on separate days in a counterbalanced study design: hypoglycemia (*hypo*), artificial euglycemia (*qu_art_*), and natural euglycemia (uni*_t_*). These conditions enable two within-subject contrasts: (1) The comparison between *hypo* and *eu_art_* isolates the effect of reduced blood glucose levels on CMR_O2_ (glucose contrast), as both conditions maintained equivalent insulin levels (Fig. 1A). (2) The contrast between *eu_art_* and *eu_nat_* examines the effect of hyperinsulinemia (insulin contrast), with euglycemia maintained in both conditions (Fig. 1A). Additionally, we investigated whether prior hypoglycemia affects cognitive performance even after glucose levels have been restored to euglycemia. Therefore, participants completed cognitive tests assessing memory and attention following the MR scan once euglycemia was re-established. A schematic timeline of the study setup is provided in Fig. 1B.

**Fig. 1.**
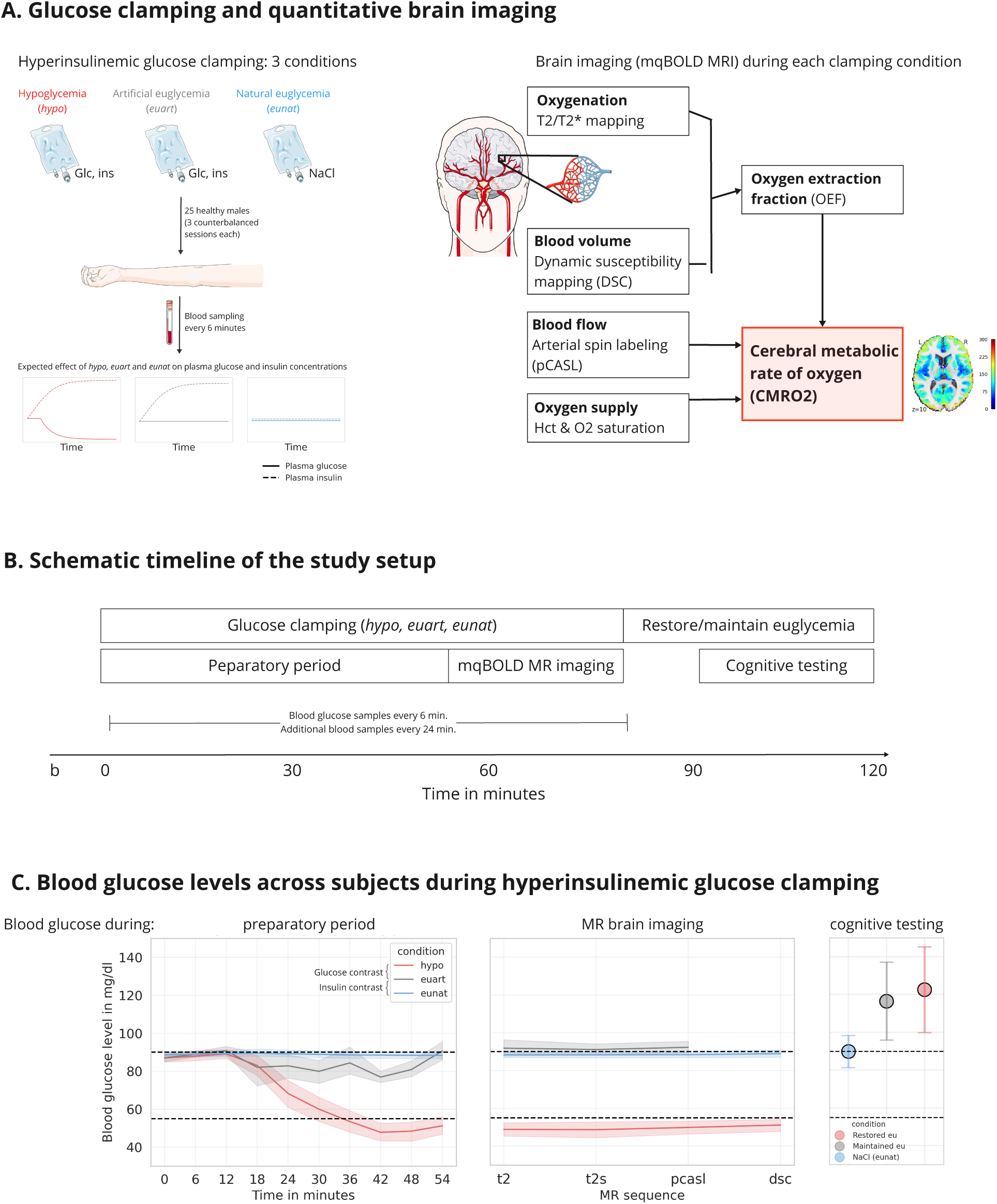
Study design and data processing. On three different days, we induced hypoglycemia (*hypo*; glucose, insulin), artificial euglycemia (*eu_art_*; glucose, insulin) and natural euglycemia (*eu_nat_*; NaCl) in healthy male subjects in a counterbalanced, single-blind within-subjects design. n_hypo_=23, n_euart_=19, n_eunat_=25. **A. Hyperinsulinemic clamping and mqBOLD MR imaging.** *Left:* To reach the three conditions (*hypo, eu_art_, eu_nat_*), continuous infusions of glucose and insulin or NaCl were administered, with blood glucose monitored at 6-minute intervals. Once blood glucose targets were reached, brain imaging was performed while infusion and monitoring continued. *Right:* Multiparametric quantitative (mqBOLD) brain imaging provides voxelwise CMR_O2_ values based on deoxygenation, cerebral blood flow and cerebral blood volume parameters. Figure adapted from (Hechler et al., 2023). **B. Schematic timeline of the study setup.** After baseline measurements (t=b), hyperinsulinemic glucose clamping was initiated at t=0 to induce *hypo* or *eu_art_*. In *eu_nat_*, only NaCl was infused. Once blood glucose levels stabilized, MR scanning was performed (t≈55), while glucose clamping continued. Following the MR scan, euglycemia was either restored (after prior *hypo*) or maintained (after prior *eu_art_*). Finally, we conducted cognitive testing. Independent of the blood glucose levels during MR scanning, participants were always euglycemic during the period of cognitive testing. **C. Blood glucose levels across subjects during hyperinsulinemic glucose clamping.** Shown are means and standard deviation. Black dashed lines indicate glucose targets (euglycemia: 90 mg/dL and hypoglycemia: 55 mg/dL). *Left:* Blood glucose levels during the preparatory phase before MR scanning, with clamping initiated at t=0, MR scanning at t≈55. *Middle:* Blood glucose levels during MR scanning for each MR imaging sequence, calculated as the average measurements taken before and after each sequence. *Right:* Blood glucose levels during cognitive testing. Here, euglycemia was either restored (after prior *hypo*) or maintained (after prior *eu_art_*). There were no significant differences in blood glucose levels between *restored eu* and *maintained eu* during cognitive testing.

Subjects presented with average fasting blood glucose levels of 87.33 mg/dL (sd=4.8). To verify stable glucose metabolism, tissue glucose levels were monitored for seven days via a continuous glucose monitoring system (FreeStyle Libre 2, Abbott Laboratories) prior to study participation (average=86.55 mg/dL, sd=7.53). During MR scanning, subjects’ average blood glucose levels were 49.66 mg/dL (sd=8.92) during *hypo*, 91.75 mg/dL (sd=7.95) during *eu_art_* and 88.51 mg/dL (sd=4.44) during *eu_nat_* (Fig. 1C).

Additionally, during the MR scan, insulin levels were significantly different between insulin contrast conditions (*eu_art_* vs. *eu_nat_*; Wilcoxon signed-rank test: W=0, p<0.001; Fig. S5). Importantly, insulin levels did not differ between *hypo* and *eu_art_* (Wilcoxon signed-rank test: W=61.0, p=0.49; Fig. S5), ensuring a pure glucose contrast. As expected, hypoglycemia significantly increased epinephrine levels (Wilcoxon signed-rank test: W=3.0, p<0.001; Fig. S3) and significantly decreased c-peptide levels (Wilcoxon signed-rank test: W=9.0, p<0.01; Fig. S2) compared to *eu_art_*. Furthermore, no differences in oddball task performance during the MR scan were observed across conditions (Friedman’s test: *χ*^2^_F_=0.49, p=0.78), indicating similar levels of wakefulness in all conditions. Aside from the changes reported here, no significant differences in blood parameters were observed between conditions during the MR scan. More information on subjects’ physiological characteristics, as well as all blood and statistical parameters, can be found in the supplementary material (Tables S1, S3, Fig. S1-S6).

### mqBOLD imaging analysis

#### Neither hypoglycemia nor hyperinsulinemia affect CMR_O2_

We analyzed the imaging data using a linear mixed effects model to examine the impact of condition on CMR_O2_ across different functional brain networks (Table 1). Our model, CMR_O2_ ∼ condition*network + (1|subject/condition), tested the effect of conditions (*hypo, eu_art_, eu_nat_*) on CMR_O2_ in each of seven functional brain networks defined by Yeo et al. (2011), while accounting for the within-subjects design. To reduce voxel-level noise, CMR_O2_ input data were provided as gray matter voxel medians per region of interest (ROI) using an established cortical atlas (Schaefer et al., 2018), which defines functionally homogenous ROIs across 400 regions and differentiates large-scale sensory from higher-order cognitive systems. The cortical distribution of CMR_O2_ across networks and regions per condition is depicted in Fig. 2A.

**Fig. 2.**
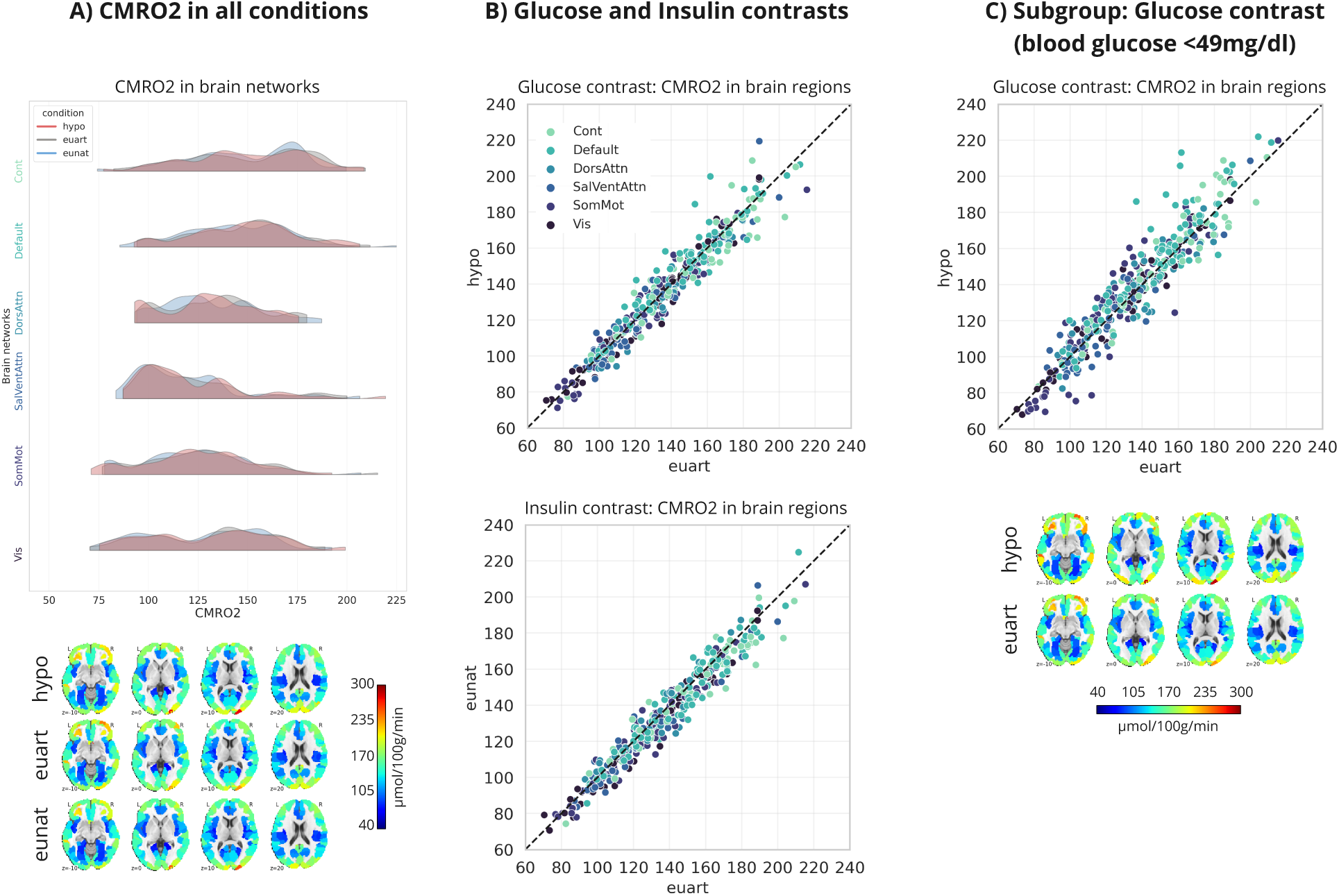
Neither hypoglycemia nor hyperinsulinemia affect CMR_O2_. For all tests and plots, contrasts included the glucose contrast (*hypo* vs. *eu_art_*) and the insulin contrast (*eu_nat_* vs. *eu_art_*). Significant levels: <0.001: ***, <0.01: **, <0.05: *. Network abbreviations: Cont ≙ Control; Default ≙ Default mode; DorsAttn ≙ Dorsal attention; SalVentAttn ≙ Salience; SomMot ≙ Somatomotor. All analyses were performed in native space. **A. CMR_O2_ in all conditions.** The ridgeline plots descriptively show the distribution of regional CMR_O2_ values, averaged across subjects, per condition and functional brain network. These regional CMR_O2_ values are also displayed in the axial brain slices below. **B. Glucose and insulin contrasts.** The regression plots descriptively show the similarity between regional CMR_O2_ values during *hypo* and *eu_art_* (glucose contrast; top) and *eu_nat_* and *eu_art_* (insulin contrast; bottom). The linear mixed effects model did not reveal significant differences between conditions in any brain network (see Results section). Datapoints reflect regional parameter values, averaged across subjects, color-coded by functional brain network. The dashed line is the angle bisector. **C. Subgroup: Glucose contrast.** Similar to the main group, in the more severely hypoglycemic subgroup (n=6), the linear mixed effects model did not reveal any CMR_O2_ differences in any network. CMR_O2_ levels were thus kept stable even under more severe hypoglycemia.

**Table 1.**
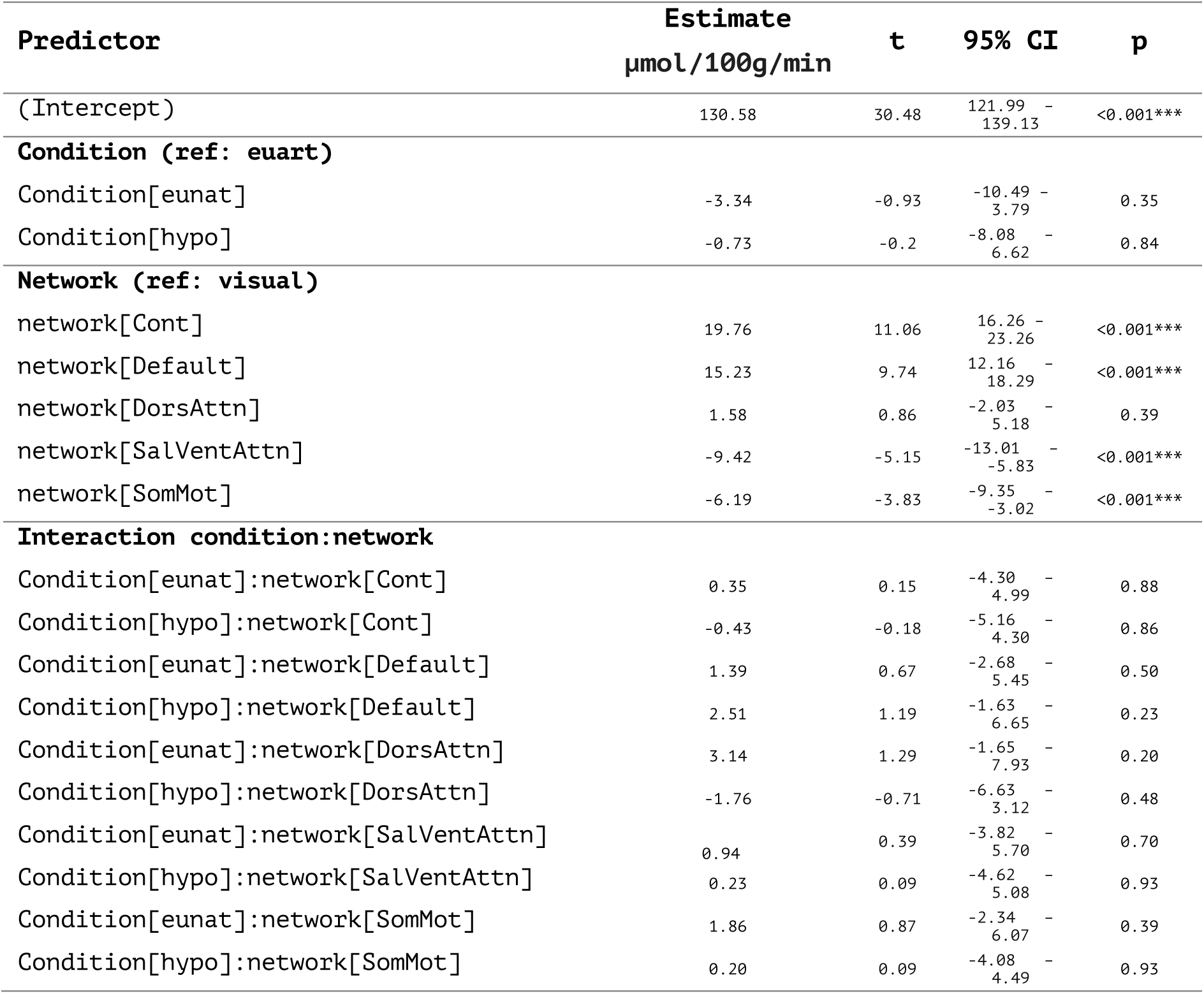
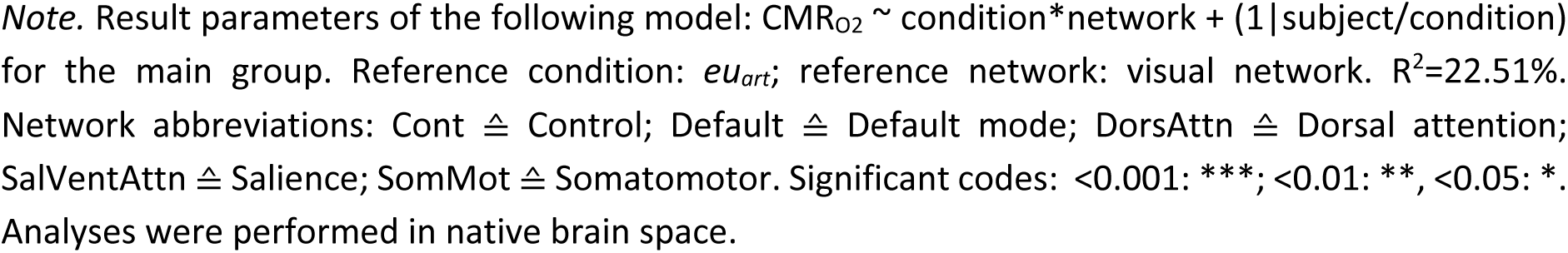
Results of the linear model predicting CMR_O2_ in the main group.

Initially, we compared our model to a null model, excluding the condition term, to determine whether the intervention significantly improved model fit. Including the condition term accounted for significantly more variance in CMR_O2_ (χ^2^=865.88, p<0.001), with the combined fixed and random effects explaining 22.51% of the variance. We also tested whether including epinephrine as an additional interaction term would enhance model fit, given its significant variation between conditions during MR scanning. However, this did not yield a significant improvement (χ^2^=25.26, p=0.24).

Based on the model’s intercept, the estimated baseline CMR_O2_ for participants in the reference condition (*eu_art_*) and reference network (visual network) was 130.58 μmol/100g/min (Table 1). The main effects of *condition* were not significant, as neither *eu_nat_* nor *hypo* showed a significant difference in CMR_O2_ compared to the reference condition (*eu_art_*) within the reference network (estimate: -3.34, t=-0.93, p=0.35 for *eu_nat_*; estimate: -0.73, t=-0.2, p=0.84 for *hypo*). The estimates in the model output can be interpreted neurobiologically as absolute deviations from the intercept, representing the change in CMR_O2_ relative to the baseline value in the reference condition and network in μmol/100g/min. The main effects of *network* did show significant differences in CMR_O2_ during the *eu_art_* condition: There were differences between the visual network and the control network (estimate: 19.76, t=11.06, p<0.001), the default mode network (estimate: 15.23, t=9.74, p<0.001), the salience network (estimate: -0.42, t=-5.15, p<0.001) and the somatomotor network (estimate: -6.19, t=-3.83, p<0.001). These network differences, observed within condition, highlight the spatial variability of CMR_O2_ across brain networks (Fig. 2A, ridgeline plots) and brain regions (Fig. 2A, brain slices). Specifically, higher cognitive networks, such as the control and default mode networks, were the energetically most expensive networks (150.34 μmol/100g/min and 145.81 μmol/100g/min, respectively), followed by the dorsal attention (132.16 μmol/100g/min) and visual networks (130.58 μmol/100g/min). The networks with the lowest CMR_O2_ were the somatomotor (124.39 μmol/100g/min) and salience networks (121.16 μmol/100g/min). However, the main effects do not provide insight into how network differences vary between conditions, which is addressed by the interaction effects.

The interaction term directly tests the key research question of this paper: whether oxygen metabolism in different functional brain networks varies depending on the glucose (*hypo* vs. *eu_art_*) or insulin (*eu_nat_* vs. *eu_art_*) contrasts, or both. Specifically, the *condition[cond]:network[nw]* interaction tests whether the effect of the condition *cond* on CMR_O2_ in network *nw* differs significantly from the effect of the condition *eu_art_* on CMR_O2_ in the visual network. In our case, none of the interaction terms is significant (Table 1), indicating that neither hypoglycemia nor hyperinsulinemia had a significant effect on CMR_O2_ in any of the functional brain networks. Thus, under both glucose deficiency and insulin prevalence, cortical oxygen metabolism is kept at a normal level (Fig. 2B, regression plots).

Next, we were interested in whether the robustness of CMR_O2_ observed under mild hypoglycemia would uphold even under more severe hypoglycemia. To address this, we defined a subgroup of participants within our study cohort who experienced more severe hypoglycemia, based on previously established glycemic thresholds for neuroglycopenic symptoms (Mitrakou et al., 1991). Accordingly, this subgroup included participants whose blood glucose levels remained consistently below 49 mg/dL during MR scanning, comprising a total of six participants in total. Similar to the main group, this subgroup did not exhibit differences in CMR_O2_ in response to hypoglycemia (Fig. 2C, Table S5). This suggests that, even under severe glucose deficiency, cortical oxygen metabolism rates remain unaffected.

#### Hypoglycemia leads to widespread CBF increases

As a next step, we were interested in whether hypoglycemia and hyperinsulinemia did not affect cerebral metabolism at all or whether solely CMR_O2_ was spared. Therefore, we investigated the effect of our interventions on CBF as a subcomponent of CMR_O2_. The cortical distribution of CBF across networks and regions per condition is depicted in Fig. 3A and 3B. We used the same linear model as previously described: CBF ∼ condition*network + (1|subject/condition), which explained 44.7% of the variance in CBF. We found a significant increase in CBF under *hypo* compared to *eu_art_* in the control network (estimate: 2.29, t=3.99, p<0.001), default mode network (estimate: 1.95, t=3.84, p<0.001) and salience network (estimate: 1.20; t=2.02, p<0.05) (Tables 2, S6 and Fig. 3A, 3C). Again, the estimates can be interpreted in ml/100g/min as the deviation from the intercept (43.56 ml/100g/min). Notably, the higher cognitive networks, specifically the control and default mode networks, showed the most pronounced increases in CBF under hypoglycemia. These networks also exhibited the highest baseline metabolic rates (Table 1). Additional analyses showed that baseline CMR_O2_ and CBF increase under hypoglycemia correlated significantly (r=0.23, p<0.001; Fig. S7), suggesting a relationship between energy metabolism and CBF increase under hypoglycemic conditions.

**Fig. 3.**
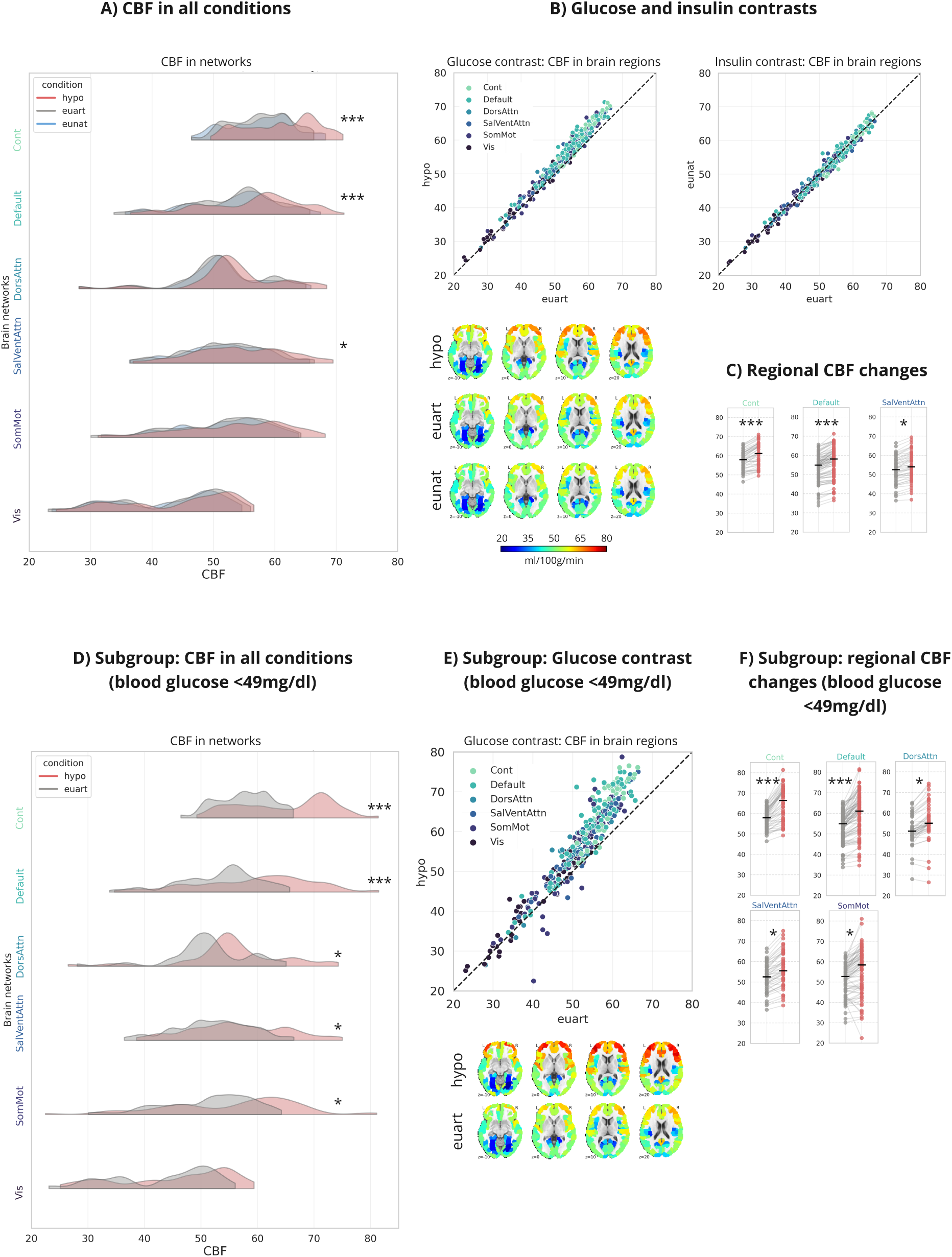
Hypoglycemia increases CBF, hyperinsulinemia has no effect. For all tests and plots, contrasts included the glucose contrast (*hypo* vs. *eu_art_*) and the insulin contrast (*eu_nat_* vs. *eu_art_*). Significant levels: <0.001: ***, <0.01: **, <0.05: *. Network abbreviations: Cont ≙ Control; Default ≙ Default mode; DorsAttn ≙ Dorsal attention; SalVentAttn ≙ Salience; SomMot ≙ Somatomotor. All analyses were performed in native space. **A. CBF in all conditions.** The ridgeline plots descriptively show the distribution of regional CBF values, averaged across subjects, per condition and functional brain network. The linear mixed effects model revealed significant CBF increases during *hypo* compared to *eu_art_* in the control, default mode and salience networks. **B. Glucose and insulin contrasts.** The regression plots descriptively show CBF differences between *hypo* and *eu_art_* (glucose contrast; left) and *eu_nat_* and *eu_art_* (insulin contrast; right). Datapoints reflect regional parameter values, averaged across subjects, color-coded by functional brain network. The dashed line is the angle bisector. Regional CBF values are also displayed in the axial brain slices below. **C. Regional CBF changes.** The paired point plots show the change of each brain region between *eu_art_* and *hypo* for the significant brain networks, with black bars illustrating the median value. **D. Subgroup: CBF in all conditions.** The ridgeline plots descriptively show the distribution of regional CBF values, averaged across subjects in the subgroup (n=6), per condition and functional brain network. This subgroup included participants with more severe hypoglycemia (blood glucose levels of <49mg/dl). The more severe hypoglycemia amplified the effect observed in the main group: The linear mixed effects model revealed significant CBF increases during *hypo* compared to *eu_art_* in the control, default mode, dorsal attention, salience and somatomotor networks. **E. Subgroup: Glucose contrast.** The regression plots descriptively show CBF differences between *hypo* and *eu_art_* (glucose contrast) in the subgroup. The subgroup comprises the same individuals as described in D. Datapoints reflect regional parameter values, averaged across subjects, color-coded by functional brain network. Regional CBF values are also displayed in the axial brain slices below. **F. Subgroup: Regional CBF changes.** The paired point plots show the change of each brain region between *eu_art_* and *hypo* for the significant brain networks in the subgroup, with black bars illustrating the median value.

**Table 2.**
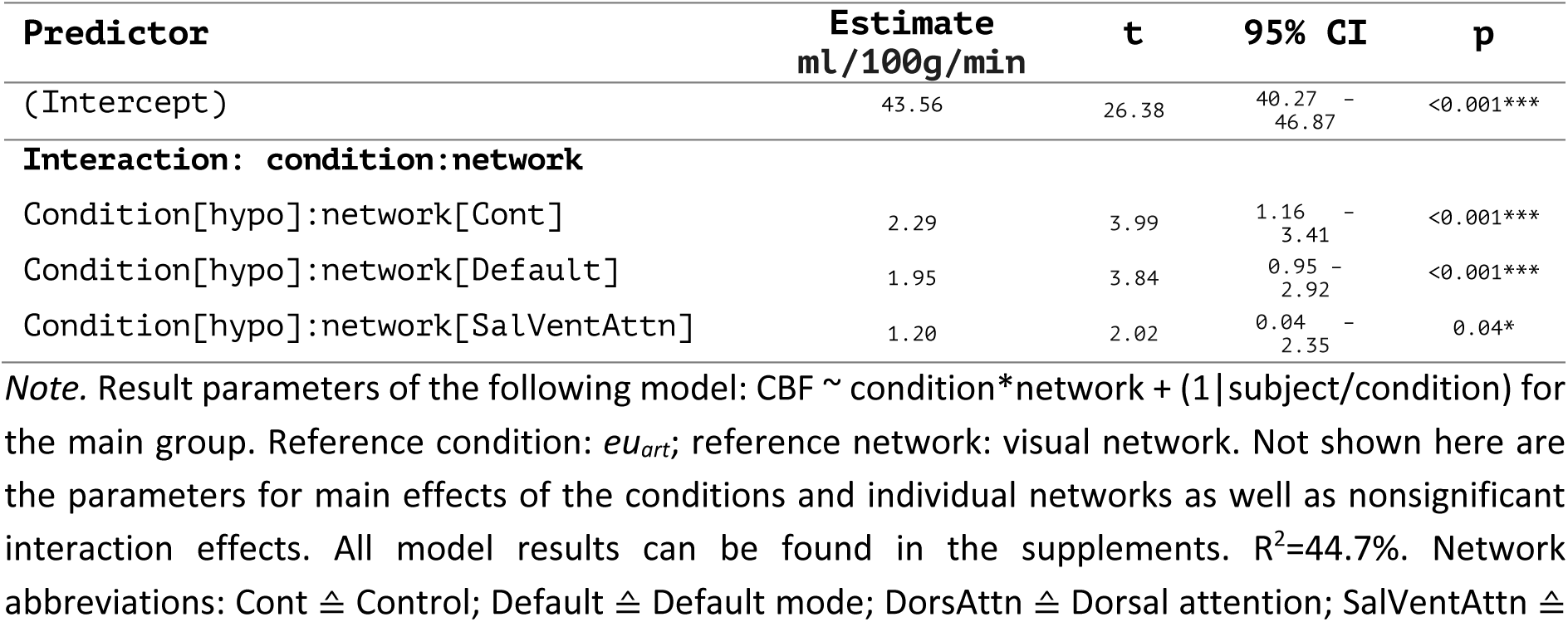
Results of the linear model predicting CBF in the main group.

| Predictor | Estimate<br>ml/100g/min | t | 95% CI | p |
| --- | --- | --- | --- | --- |
| (Intercept) | 43.56 | 26.38 | 40.27 - 46.87 | <0.001*** |
| **Interaction: condition:network** |  |  |  |  |
| Condition[hypo]:network[Cont] | 2.29 | 3.99 | 1.16 - 3.41 | <0.001*** |
| Condition[hypo]:network[Default] | 1.95 | 3.84 | 0.95 - 2.92 | <0.001*** |
| Condition[hypo]:network[SalVentAttn] | 1.20 | 2.02 | 0.04 - 2.35 | 0.04* |

*Note.* Result parameters of the following model: $CBF \sim \text{condition} * \text{network} + (1 | \text{subject/condition})$ for the main group. Reference condition: *eu<sub>art</sub>*; reference network: visual network. Not shown here are the parameters for main effects of the conditions and individual networks as well as nonsignificant interaction effects. All model results can be found in the supplements. $R^2=44.7\%$ . Network abbreviations: Cont $\triangleq$ Control; Default $\triangleq$ Default mode; DorsAttn $\triangleq$ Dorsal attention; SalVentAttn $\triangleq$

In contrast, hyperinsulinemia did not affect CBF, as can be derived from the non-significant interaction terms for the *eu_nat_* condition (Table S6, Fig. 3B right regression plot). This indicates that insulin has no significant effect on cerebral hemodynamics or oxygen metabolism.

In the severely hypoglycemic subgroup, CBF changes were more prominent than in the main group (Fig. 3D-F). The intercept was at 43.08 ml/100g/min. We found significantly increased CBF rates in the control network (estimate: 4.79, t=5.74, p<0.001), default mode network (estimate: 3.89, t=5.32, p<0.001), dorsal attention network (estimate: 2.03, t=2.35, p<0.05), salience network (estimate: 1.93, t=2.25, p<0.05) and somatomotor network (estimate: 1.63, t=2.15, p<0.05) (Tables 3, S7 and Fig. 3D, 3F). Hence, CBF increased in every network but the visual network in response to more severe hypoglycemia.

**Table 3.**
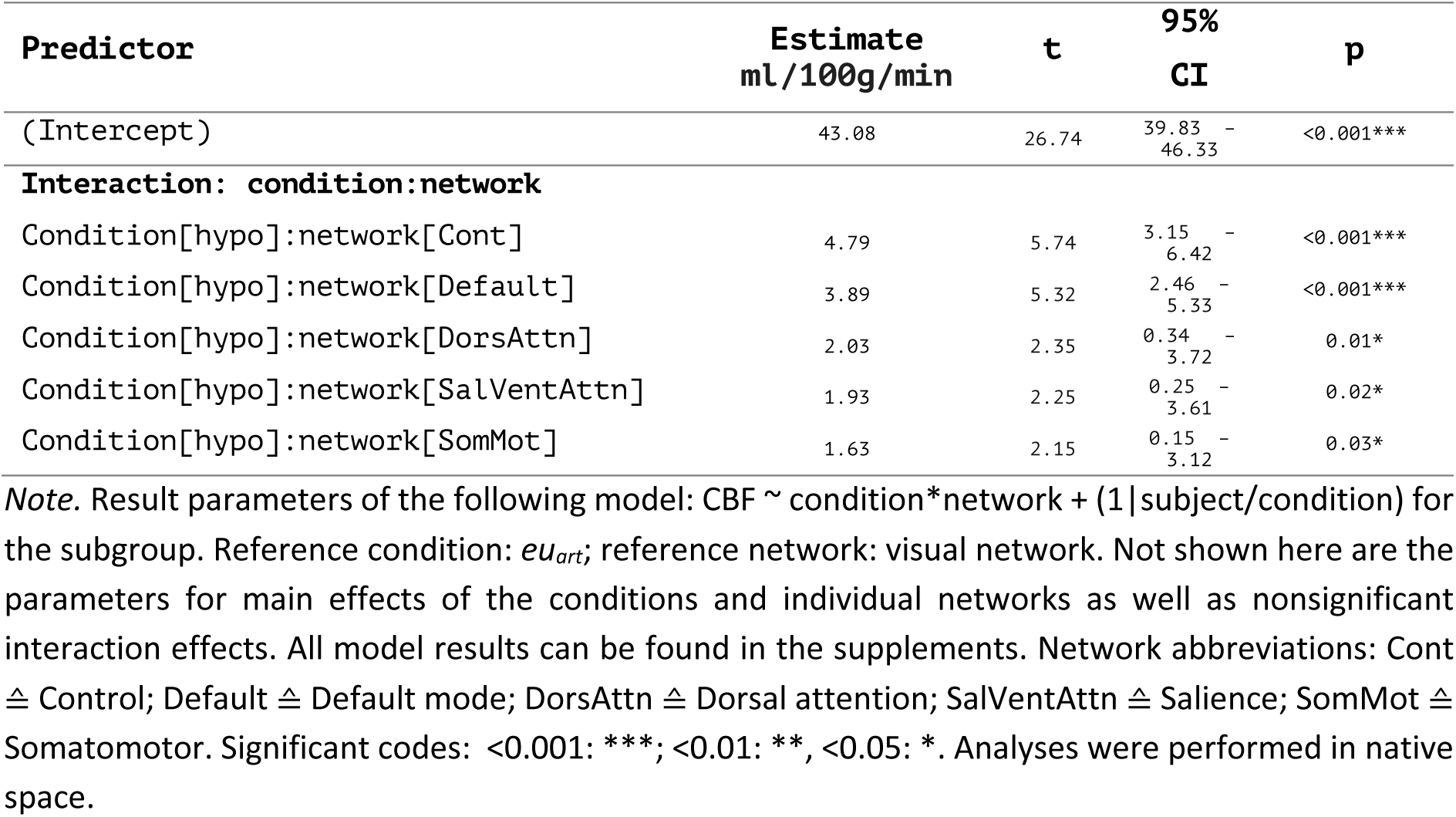
Results of the linear model predicting CBF in the subgroup.

### Behavioral analysis

Behavioral data were acquired after the MR scan, once euglycemia was *restored* (after prior *hypo* (blood glucose mean ± sd = 125.17 mg/dL ± 26.27) or *maintained* after prior *eu_art_* (blood glucose mean ± sd = 117.38 mg/dL ± 21.48). The restoration of euglycemic levels after *hypo* was successful as indicated by non-significant differences between prior *hypo* and prior *eu_art_* (Wilcoxon signed-rank test: W=108.5, p=0.39). For clarity, we will refer to these conditions as *restored euglycemia* (prior *hypo*) and *maintained euglycemia* (prior *eu_art_*), respectively. At the point of cognitive testing, there were no significant differences in epinephrine levels (Friedman’s test; *χ*^2^_F_=4.79, p=0.09) or any other stress parameter between the conditions. Further, the previously observed reductions in c-peptide in response to hypoglycemia were no longer present once euglycemia was restored (restored vs. maintained euglycemia: Wilcoxon signed-rank test; W=36.0, p=0.3). These findings indicate that not only were blood glucose levels normalized, but physiological responses to prior hypoglycemia had also subsided.

#### Prior hypoglycemia impairs memory consolidation but does not affect learning

In the memory task, participants were instructed to learn a list of 15 concrete nouns presented sequentially on a screen and were then asked to recall as many of the learned nouns as possible. Subjects did not show differences between restored and maintained euglycemia in memory encoding performance (Friedman’s test; *χ*^2^_F_=1.37, p=0.5; Fig. 4A) or in memory recall after a 20-minute consolidation period (Friedman’s test; *χ*^2^_F_=2.63, p=0.27; Fig. 4B). However, memory recall after a 24-hour consolidation period was significantly impaired in the restored vs. maintained euglycemia condition (Wilcoxon signed-rank test; W=14.0, p<0.01), even after controlling for initial learning performance (GLM, p=0.03 for the effect of restored vs. maintained euglycemia on memory consolidation 24 hours later) (see Fig. 4C). Including both condition and learning performance as predictors explained 60.7% of the variance in memory recall after 24 hours. There were no differences in memory performance for the insulin contrast.

**Fig. 4.**
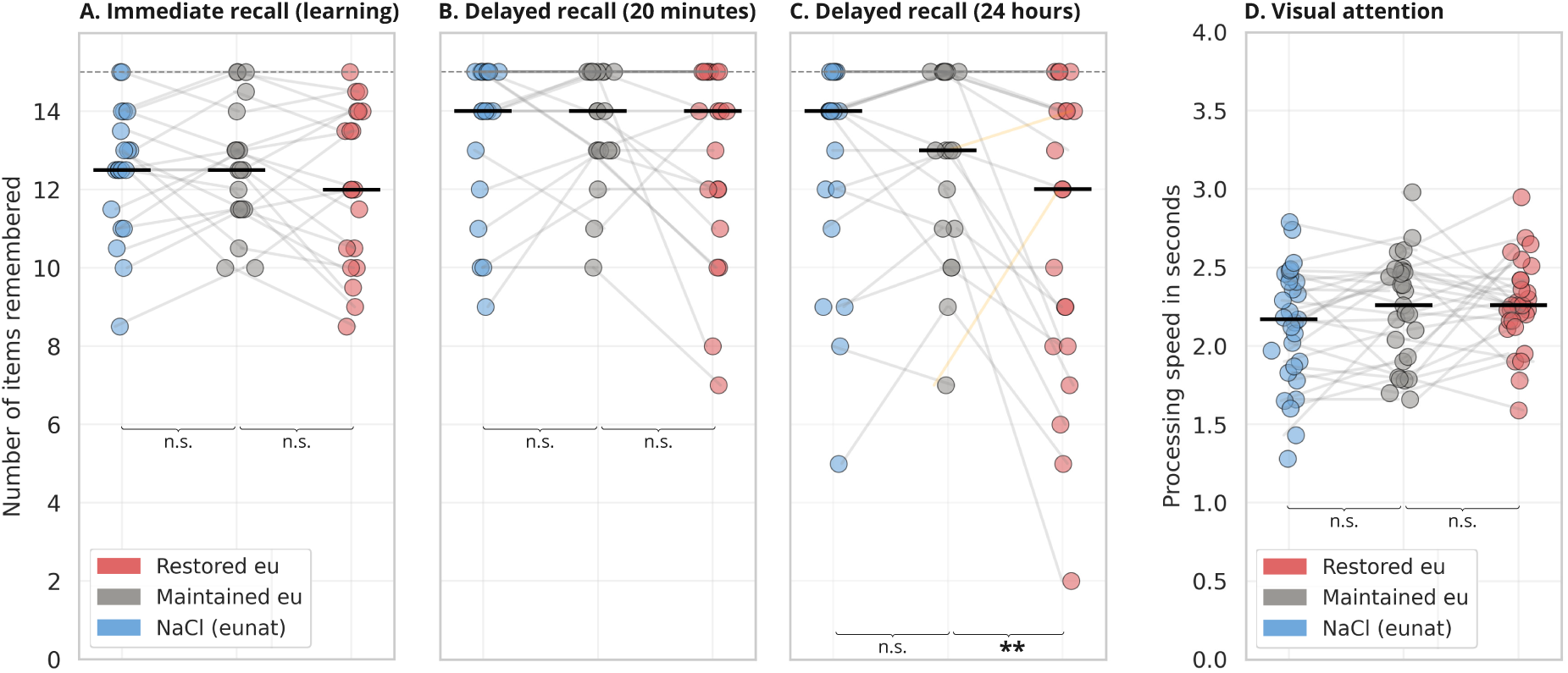
Cognitive results after glucose clamping. Cognitive tests were conducted under euglycemic conditions after prior *hypo* (*restored euglycemia*) or after prior *eu_art_* (*maintained euglycemia*). Each dot represents a subject’s performance on the respective task under the respective condition. The black bars mark the median of each sample. Significant levels: <0.001: ***, <0.01: **, <0.05: *. **A-C. Memory task results.** In the memory task, subjects were instructed to learn 15 concrete nouns (marked as dashed line at y=15) presented twice and completed recall tests: immediately after learning (averaged across both learning periods) (A), after 20 minutes (B) and after 24 hours (C). After 24 hours, recall was significantly impaired when subjects experienced hypoglycemia prior to learning, despite being euglycemic during the task. This result was upheld when controlling for initial learning level. The two orange lines represent the subjects showing an effect of opposite direction. **D. Visual attention task results.** Subjects identified a target stimulus among distractor stimuli. Previous glucose status did not affect visual attention.

#### Attention returns to normal levels with restored euglycemia

To control for potential effects of attention deficits, subjects performed a visual attention task designed to measure their speed in identifying a target stimulus among similar distractors. Statistical analysis showed no significant differences in reaction times between conditions (repeated measures ANOVA; F=2.27, p=0.11), indicating that attention deficits did not impact task performance (Fig. 4D).

## Discussion

In this study, we investigated the effects of insulin-induced hypoglycemia as well as of hyperinsulinemia on cerebral oxygen metabolism (CMR_O2_) and the effect of hypoglycemia on cognitive function, particularly memory performance. In line with the Selfish Brain Theory (Peters et al., 2004), our findings indicate that CMR_O2_ remains stable during both hypoglycemia and hyperinsulinemia, highlighting the brain’s resilience in preserving its metabolic activity despite fluctuations in peripheral energy substrate levels. Notably, we found widespread CBF increases in response to hypoglycemia, particularly in higher cognitive brain networks (Thomas Yeo et al., 2011), which may reflect compensatory vascular responses to sustain cerebral energy supply under hypoglycemia. Importantly, we also found that while learning and attention remained unaffected, restored euglycemia following hypoglycemia selectively impaired long-term memory performance, pointing to potential memory-related vulnerabilities following acute glucose shortages.

Furthermore, our setup was validated by expected physiological responses: Hypoglycemia increased epinephrine levels, consistent with stress hormone elevation during low glucose conditions to stimulate gluconeogenesis (Cryer, 1993), and decreased c-peptide levels, indicating a reduction in endogenous insulin production (Woods et al., 1974). These changes confirm the effectiveness of our hypoglycemic intervention.

### Hypoglycemia leads to widespread increases in CBF but does not affect CMR_O2_

Our primary aim was to determine whether hypoglycemia induces reductions in cerebral oxygen metabolism, similar to its effects on cerebral glucose metabolism (CMR_glc_) (Blazey & Raichle, 2019). Previous studies in animals (Richardson et al., 1985) and humans (Lubow et al., 2006) have suggested varying impacts of hypoglycemia on brain metabolism but direct measurements of CMR_O2_ under hypoglycemia are lacking. fMRI, the gold standard for assessing brain activity, has been used to study the blood oxygen level dependent (BOLD) contrast during hypoglycemia, revealing a lower BOLD signal under hypoglycemia compared to euglycemia during a visual task (Anderson et al., 2006). However, the BOLD signal is a compound signal, reflecting information about both hemodynamic and metabolic changes. Therefore, it can be influenced by either increased oxygen consumption or increased CBF, making it an indirect measure of cerebral oxygen metabolism. To address these limitations, we measured CMR_O2_ via multiparametric quantitative BOLD (mqBOLD) (Christen et al., 2012; Hirsch et al., 2014; Kaczmarz et al., 2020) in the present study. By acquiring separate data on oxygenation and CBF, we were able to distinguish between purely hemodynamic processes and actual oxygen consumption.

If cerebral energy metabolism generally decreased in response to hypoglycemia, we would expect reductions in CMR_O2_ similar to those observed in CMR_glc_ (Blazey & Raichle, 2019). Nevertheless, our findings showed that CMR_O2_ remained stable even under more severe hypoglycemia, suggesting that the brain is capable of maintaining its energy metabolism, and likely related ATP production, despite reduced systemic glucose availability. This stability likely reflects the utilization of alternative energy substrates, such as astrocytic glycogen and lactate which still require oxidation for ATP production (Brown & Ransom, 2007). An alternative explanation is that CMRglc does not decrease as much as previously thought, allowing glucose to remain the primary substrate for oxidation and ATP synthesis in the brain. Since our data did not include concurrent CMRglc measurements, we cannot definitively rule out this possibility. Nevertheless, both scenarios – whether the brain shifts to alternate fuels or channels remaining glucose – demonstrate the brain’s metabolic resilience. In either case, stable CMR_O2_ levels imply that the brain sustains relatively constant ATP production at rest, highlighting its capacity to preserve normal metabolic activity even under conditions of energy substrate deficiency.

Despite no changes in CMR_O2_ in response to hypoglycemia, we observed alterations in CBF, a subcomponent of imaging-derived CMR_O2_. The fact that we observed CBF changes without corresponding CMR_O2_ changes further supports our choice of measuring CMR_O2_ via mqBOLD rather than relying on more indirect methods like conventional BOLD imaging. During hypoglycemia, there were significant increases in CBF in the control, default mode, and salience networks. This aligns with existing literature, which reports hypoglycemia-induced CBF increases in regions like the medial prefrontal cortex (Teves et al., 2004), anterior cingulate cortex (Dunn et al., 2018; Teh et al., 2010), dorsolateral prefrontal cortex, and angular gyrus (Dunn et al., 2018), all of which are regions within the three aforementioned functional networks. In contrast, animal studies report whole-brain increases in CBF in response to hypoglycemia in rats (Bryan et al., 1987; Choi et al., 2001), likely due to the lower hypoglycemia targets chosen for animal studies (∼30 mg/dL). In line with this, in our more severely hypoglycemic subgroup, the CBF effect was amplified, with significant CBF increases in every network except the visual network. Together, our findings extend beyond individual regions, highlighting the compensatory effects on brain networks involved in higher cognitive functions.

In the main group, the more pronounced increases in the higher cognitive networks (control and default mode networks) may reflect their higher metabolic demand. As demonstrated in Fig. S7, CBF increases during hypoglycemia correlate significantly and positively with baseline CMR_O2_ values (p<0.001). In general, higher rates of CBF could serve to provide the brain more efficiently with the remaining glucose and alternative substrates. Previous studies have interpreted CBF increases during hypoglycemia as markers of heightened neuronal activity in these regions, suggesting their involvement in the autonomic response to hypoglycemia (Teves et al., 2004). However, if CBF increases were due to neuronal activation, we would expect corresponding CMR_O2_ changes. Instead, CBF and CMR_O2_ appear uncoupled during hypoglycemia. Alternatively, CBF alterations could be driven by increased epinephrine levels (J. J. Lee et al., 2017; Thomas et al., 1997). In summary, we interpret the uncoupling of CMR_O2_ and CBF changes, along with the more pronounced CBF increases in higher cognitive brain networks, as a compensatory mechanism aimed at supplying these energy-demanding regions with adequate substrates.

### Hyperinsulinemia does not affect brain oxygen metabolism

The precise role of insulin in the brain is not completely understood. Insulin functions not only as a signaling molecule within the central nervous system but also plays a role in systemic energy homeostasis, influencing both glucose sensing and regulation (García-Cáceres et al., 2016; Herrera Moro Chao et al., 2022). Although cerebral glucose uptake largely occurs independently of insulin, facilitated primarily through the glucose transporters GLUT1 and GLUT3, insulin-sensitive glucose transporter GLUT4 is also widely expressed in various brain regions, with particularly high concentrations in the hippocampus (El Messari et al., 2002; Koepsell, 2020). This raises the question of whether elevated insulin levels lead to increased cerebral glucose metabolism, as is the case in the periphery. Evidence suggests that cerebral glucose metabolism remains unchanged during hyperinsulinemia in healthy individuals (Hirvonen et al., 2011), supporting the notion that insulin may not significantly influence glucose uptake in the brain. In line with these findings, our results showed no changes in CMR_O2_ across the entire cortex under hyperinsulinemia.

The effect of insulin on cerebral blood flow (CBF) is less clear, with studies reporting conflicting results. While some studies have found stable CBF levels during hyperinsulinemia (Hirvonen et al., 2011), others report reductions in CBF with elevated insulin (Kullmann et al., 2015). These differences might reflect a more complex interaction, potentially modulated by factors such as age (Akintola et al., 2017) or peripheral insulin sensitivity (Kullmann et al., 2015). In our study, we observed no significant effects of hyperinsulinemia on CBF, indicating that insulin may not substantially alter CBF in the young, healthy population.

### Cognition

In addition to studying cerebral oxygen dynamics, the present study was conducted to investigate whether prior hypoglycemia, once restored to euglycemia, still affects cognition. Previous studies have shown memory deficits during acute episodes of hypoglycemia (Sommerfield et al., 2003b, 2003a). However, interpreting these results is complicated by the concurrent attention deficits observed during hypoglycemia (Ewing et al., 1998; Graveling et al., 2013b; McAulay et al., 2001b), as reduced attentiveness can contribute to poorer performance on cognitive tasks, making it difficult to disentangle the specific effects of hypoglycemia on memory alone.

Our results suggest that while attention and encoding are unaffected, restored euglycemia after a hypoglycemic period impairs memory consolidation. The unaltered processing speed in the attention task suggests that (a) attention deficits appear to be restricted to periods of acute hypoglycemia and (b) impaired consolidation during restored euglycemia seems to reflect a domain-specific issue rather than a general cognitive impairment. One could speculate whether the clear difference between attention and memory performance post-hypoglycemia could indicate an uncoupling of these processes during acute hypoglycemia as well. Further, the fact that encoding did not differ between conditions rules out the possibility that impaired consolidation is mediated by poorer learning ability during restored euglycemia compared to maintained euglycemia.

The underlying mechanisms of decreased memory consolidation when learning under restored euglycemia are not entirely clear yet in line with an energy-demanding consolidation process (Sgammeglia & Sprecher, 2022). It is well established that there is an increased energy demand during long-term memory formation in the hippocampus (El Messari et al., 2002; McNay et al., 2000). The decreased glucose metabolism during low blood glucose levels might thus account for memory deficits during acute hypoglycemia but it cannot account for the impairment of memory consolidation specifically during restored euglycemia, especially with learning remaining unaffected. In this context, past studies suggest that lactate shuttling from glial cells to neurons plays an important role in long-term, but not short-term, memory (Gao et al., 2016; Newman et al., 2011; Suzuki et al., 2011). This lactate comes primarily from glycogen stored in astrocytes. Under hypoglycemia, these glycogen repertoires are depleted (Öz et al., 2007), hence leaving no glycogen for subsequent long-term memory formation. Additionally, inhibiting glycogen breakdown has been shown to impair long-term memory formation, with glucose unable to substitute for the lack of glycogen (Gibbs et al., 2006; Suzuki et al., 2011). This demonstrates that glycogen is not just a storage form of glucose but serves specific functions in memory consolidation.

Moreover, animal studies have found that about 30 minutes after learning, glycogen decreases in the forebrain, scaling with elevated levels of glutamate and glutamine, suggesting glutamate/glutamine synthesis from glycogen to promote memory consolidation (Bak et al., 2018; Hertz et al., 2003; O’Dowd et al., 1994). Supporting synaptic plasticity, glutamate is crucial for memory formation (Barnes et al., 2020). Depleted glycogen reserves due to prior hypoglycemia could thus lead to decreased de novo synthesis of glutamate after learning and, consequently, impaired memory consolidation. In addition, glutamate is essential for sharp wave ripples (SWRs) (Behrens et al., 2005; Colgin et al., 2004; Maier et al., 2003; Papatheodoropoulos & Kostopoulos, 2002), which are oscillatory patterns of neural activity in the hippocampus observed during periods of rest. They are thought to play a crucial role in long-term memory formation (Schreiner et al., 2023; Yang et al., 2024) and are even considered a cognitive biomarker for episodic memory consolidation and retrieval (Buzsáki, 2015). Thus, the depletion of astrocytic glycogen due to hypoglycemia and subsequent deficiency in glutamate-glutamine cycling could explain why we find impaired memory consolidation, specifically, while other cognitive domains remain unaffected.

### Conclusion

The present study investigated cerebral oxygen metabolism under hypoglycemia and hyperinsulinemia. Our findings indicate that hypoglycemia does not affect CMR_O2_ levels, suggesting the utilization of alternative energy substrates to maintain steady cerebral energy metabolism even under severe hypoglycemic conditions. It is important to note, however, that we did not directly measure the metabolism of glucose or alternative substrates in this study. Future studies that simultaneously measure both oxygen and glucose metabolism during hypoglycemia would provide more reliable insights into the potential uncoupling of CMR_O2_ and CMR_glc_. Hyperinsulinemia did not show any effects in the measured parameters, while hypoglycemia led to significant increases in CBF in large parts of the brain. Additionally, we examined the effect of restored euglycemia vs. maintained euglycemia on cognition. Memory consolidation was significantly impaired under restored euglycemia, while learning as well as attention remained unaffected. Overall, these findings demonstrate the brain’s flexibility in maintaining its energy metabolism by adapting to alternative substrates even during acute severe hypoglycemia. Nonetheless, our behavioral results highlight that brain function does not remain fully unimpaired under such conditions and suggest that the effects of hypoglycemia may be more enduring than previously thought.

## Methods

### Participants

A total of 38 healthy male human participants with no family history of metabolic disorders were recruited for this study, which comprised three sessions on separate days. Seven participants were excluded due to abnormal reactions to the experimental setup (e.g., signs of insulin resistance or poor vein status), and one participant dropped out after the first session. Four additional participants were excluded from imaging analyses due to insufficient MR data quality. To ensure hypoglycemia during MR scans, only participants whose blood glucose concentrations did not excel 65 mg/dL during hypoglycemic MR acquisition were included. This resulted in a final imaging sample size of 25 participants (mean age=23.96 ±2.3 years), with 23 sessions for the hypoglycemic condition, 25 for natural euglycemia and 19 for artificial euglycemia. It is important to note that high exclusion and dropout rates were anticipated due to the complexity and invasiveness of the study setup. Cognitive data were available from 20 participants (mean age=23.65 ±2.08 years).

To scan for major abnormalities in glucose metabolism, participants wore a continuous glucose monitoring device (FreeStyle Libre 2, Abbott Laboratories) for seven days. The sensor was attached to the upper arm and continuously measured interstitial glucose levels every 15 minutes via an integrated needle. The ethics board of the university hospital of the Technical University of Munich approved the experimental protocol, and all participants gave written informed consent prior to the study.

### Study procedure

On three separate days, we induced hypoglycemia (*hypo*; 55 mg/dL blood glucose concentration), artificial euglycemia (*eu_art_*; 90 mg/dL) and natural euglycemia (*eu_nat_*; ∼90 mg/dL) in each subject using a hyperinsulinemic glucose clamp technique (Heise et al., 2016). Participants were blind to the counterbalanced order of conditions. Given that insulin sensitivity decreases throughout the day (A. Lee et al., 1992; Saad et al., 2012), all data were acquired in the early morning after participants had fasted overnight. Upon arrival at the study site, intravenous catheters were placed in both arms, and baseline parameters were measured, including levels of epinephrine, norepinephrine, cortisol, IGF-1, insulin, c-peptide, CRP, creatinine, hematocrit, and arterial oxygen saturation. A baseline blood gas analysis (BGA; Epoc®, Epocal Inc) was also performed to obtain real-time blood glucose levels. Acquisition of the stress hormones epinephrine, norepinephrine and cortisol served to control for stress effects on our outcome measures, particularly since epinephrine is known to increase during hypoglycemia, triggering gluconeogenesis from the kidneys and liver (Cryer, 1993). IGF-1 was measured to control for its potential effects in glucose utilization and insulin sensitivity (Clemmons, 2004; Hernandez-Garzón et al., 2016). Insulin and c-peptide provided additional information on endogenous insulin production and synthesis. While these parameters were measured repeatedly throughout the experiment, CRP, creatinine and hematocrit were only measured at baseline level to control for effects of acute infection (CRP) (Powanda & Beisel, 2003) and to fulfill MRI protocol requirements (creatinine and hematocrit). After baseline measurements, we started the one-step hyperinsulinemic glucose clamp procedure using a 20% glucose solution. Insulin infusion rates were adjusted to 2.0 mU/kg*min based on body weight and maintained consistently within and across sessions. BGAs were performed at 6-minute intervals, and glucose infusion rates were adjusted accordingly to reach or maintain the target blood glucose levels of 55 mg/dL (hypoglycemia) and 90 mg/dL (artificial euglycemia) (Spinner et al., 2016). Every 24 minutes, additional blood samples were collected to measure epinephrine, norepinephrine, cortisol, IGF-1, insulin and c-peptide which were processed and analyzed in our in-house clinical chemistry laboratory. For the natural euglycemia, we followed the same protocol but infused only sodium chloride (NaCl) to maintain subjects’ natural euglycemia levels, which are usually around 90 mg/dL. Once blood glucose levels were stabilized within the target range, subjects were transferred into the MR scanner and started with MRI acquisition. During MR scanning, we continued with the clamping procedure, including infusions and blood sampling at 6-minute intervals.

#### Experimental design: Cognitive tasks

In the scanner, participants performed a simple oddball task to maintain wakefulness while allowing for resting-state data acquisition. During this task, participants focused on a white fixation cross on a dark background which turned red at varying intervals, ranging from 20 to 40 seconds. Participants were instructed to press a button whenever they observed the fixation cross turning red. This task is a necessary measure of control as hypoglycemia can induce neuroglycopenic symptoms, such as fatigue and drowsiness (Mitrakou et al., 1991), which in turn substantially lower cerebral energy metabolism (Madsen et al., 1991). Once the MR scan was completed, we stopped all infusions and transferred the participant to another room, where we continued glucose infusions until stable euglycemia was achieved. At that point, we started with cognitive testing, which comprised a memory and an attention task. Thus, these cognitive tasks were administered outside the scanner as well as under restored or maintained euglycemia.

For the memory task, participants were presented with a list of 15 concrete nouns, displayed successively on a screen (adapted from Sommerfield et al., 2003a). They were then instructed to write down as many of the words as they could remember immediately after presentation. This memory encoding and immediate recall process was repeated one more time. Following a 20-minute interval, participants were prompted to recall the words again without any further learning trials. Approximately 24 hours later, participants were contacted for a final recall of the previously learned words.

During the 20-minute period of initial consolidation, participants performed a visual attention task (Quirk, 2020). In this task, they were instructed to find a “*T”* embedded among “*L”*s and use the arrows keys on a keyboard to indicate the direction in which the “*T*” was rotated. Any remaining time within the 20-minute consolidation period was filled with the same undemanding oddball task as performed during the MR scan, ensuring a standardized experience for participants during the initial consolidation period. An internist was present throughout the entire glucose clamping procedure to monitor participants.

### mqBOLD acquisition parameters

In the present study, we quantified cerebral oxygen metabolism by calculating voxel-wise CMR_O2_ parameter maps using mqBOLD (Christen et al., 2012; Hirsch et al., 2014; Kaczmarz et al., 2020). This technique requires several separate MR sequences. MR data acquisition was performed on a 3T Ingenia Elition MRI scanner (Philips Healthcare, The Netherlands), using a 32-channel head coil. Each session consisted of the following MR sequences:

- Multi-echo spin-echo T2 mapping: 3D gradient spin echo (GRASE) as previously described (Kaczmarz et al., 2020). 8 echoes, TE1 = ΔTE = 6 ms, TR = 251 ms, α=90°, voxel size 2×2×3.3 mm^3^, 35 slices.
- Multi-echo gradient-echo T2* mapping: As previously described (Hirsch et al., 2014; Kaczmarz et al., 2020). 12 echoes, TE1 = 6ms, ΔTE = 5 ms, TR=2229 ms, α=30°, voxel size 2×2×3 mm^3^, gap 0.3 mm, 35 slices.
- Pseudo-continuous arterial spin labeling (pCASL): As previously described (Alsop et al., 2015). Implementation according to previous literature (Göaler et al., 2019; Kaczmarz et al., 2020). PLD: 1800 ms, label duration: 1800 ms, 4 background suppression pulses, 2D EPI readout, TE=11 ms, TR=4500 ms, α=90°, 20 slices, EPI factor: 29, acquisition voxel size: 3.28×3.50×6.00 mm^3^, gap: 0.6 mm, 30 dynamic scans, including a proton density-weighted M0 scan.
- Dynamic susceptibility contrast (DSC): As previously described (Hedderich et al., 2019). A gadolinium-based contrast agent was injected as a bolus aÄer 5 dynamic scans. Dosage: 0.2ml/kg body weight, split into two injections of 0.1ml/kg body weight for two conditions (*hypo, eu_nat_*). For the third condition (*eu_art_*), DSC data could not be acquired without exceeding the recommended dosage for healthy participants. Neither could the total dosage of 0.2ml/kg body weight be further divided without compromising signal quality. For data processing of *eu_art_*, the DSC from *eu_nat_* was used. Flow rate: 4ml/s, plus 25ml NaCl. Single-shot GRE-EPI, EPI factor: 49, 80 dynamic scans, TR=2.0s, α=60°, acquisition voxel size: 2×2×3.5 mm^3^, 35 slices. Prior to CA administration, healthy kidney function was ensured. The CA was only injected at creatinine levels of ≤ 1.2 mg/dL.
- Additionally, anatomical data was acquired in one session for anatomical reference and to exclude brain lesions. This included a T1-weighted 3D MPRAGE pre- and post-gadolinium (TI=100 ms, TR=9 ms, TE=4 ms, α=8°; 170 slices, FOV=240×252×170 mm^3^; voxel size: 1.0×1.0×1.0 mm^3^; acquisition time: 2.05 minutes) and a T2-weighted 3D fluid-attenuated inversion recovery (FLAIR) image (TR = 4800 ms; TE = 293 ms, α=40°; 140 slices; FOV=240×248.9×168 mm^3^; acquisition voxel size: 1.2×1.2×1.2 mm^3^; turbo spin-echo factor: 170; inversion delay 1650 ms; acquisition time: 2:09 minutes).

### Data processing and statistical analyses

#### mqBOLD MR data

##### CMR_O2_ calculation

The quantification of all parameter maps was performed using in-house MATLAB scripts and SPM12 (Wellcome Trust Centre for Human Neuroimaging, UCL, London, UK).

- **Cerebral blood flow (CBF)** parameter maps were derived from pCASL data by calculating the average pairwise differences of motion-corrected label and control images, along with a proton-density weighted image (Alsop et al., 2015). The resulting CBF values are expressed in ml/100g/minute.
- **R2’, the transverse, reversible relaxation rate**, was calculated using the formula:

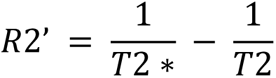

as described in previous studies (Blockley et al., 2013, 2015; Bright et al., 2019).
- **Cerebral blood volume (CBV)** parameter maps were calculated from the DSC data (Hedderich et al., 2019, p. 20; Kluge et al., 2016). When combined with R2’, the CBV maps were used to compute the subsequently yielded the **oxygen extraction fraction (OEF)** (Christen et al., 2012; Hirsch et al., 2014; Yablonskiy & Haacke, 1994) using the following formula:

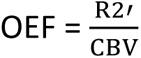
- **CMR_O2_** parameter maps were calculated by combining OEF and CBF via Fick’s principle (Fick, 1870):

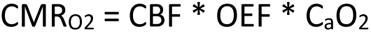

CMR_O2_ is expressed in units of μmol/100g/minute. C_a_O_2_, the arterial oxygen content, is calculated as C_a_O_2_ = 0.335*Hct*55.6*O_2_sat, where Hct is the subject’s hematocrit level and O_2_sat the arterial oxygen saturation measured with a pulse oximeter (Bright et al., 2019; Ma et al., 2020).

All individual parameter maps were registered to the first echo of the subject’s respective multi echo T2 data. We masked out the cerebellum and only considered voxels with a grey matter (GM) probability of >0.5. Additionally, in native space, we discarded voxels influenced by cerebrospinal fluid (T2>90ms), susceptibility artefacts (R2’>9s^-1^), voxels with elevated blood volume (CBV>10%, likely driven by larger vessels) and voxels with physiologically unexpected values (T2*>90ms, OEF>0.9, CBF>90).

##### Statistical analyses

For statistical analyses of the imaging data, we employed linear mixed modeling (LMM) at the network level using the R package *robustlmer* (Koller, 2016) to predict CMR_O2_. LMM was chosen due to its ability to incorporate both fixed and random effects in predicting an outcome variable. Our model included the interaction between condition and brain network (7 networks according to Thomas Yeo et al., 2011; with the limbic network excluded due to susceptibility artefacts) as a fixed effect. We set *eu_art_* as a reference condition to enable both glucose (*hypo* vs. *eu_art_*) and insulin (*eu_nat_* vs. *eu_art_*) contrasting. Further, we specified the visual network as a reference for the network variable due to previous findings suggesting it is least affected by hypoglycemia (Blazey & Raichle, 2019). The random model term (1|subject/condition) treats conditions as nested within subjects and was included to account for the repeated measures design of the study. For the dependent variable, regional CMR_O2_ values were calculated as the median voxel value within 400 Schaefer regions of interest (ROIs) (Schaefer et al., 2018) to reduce voxel-level noise. These medians were then entered as a predictor variable per subject and condition which yielded a total of 26,310 entries for the model. This resulted in the following LMM:

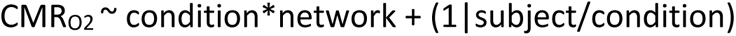

For ROI and network parcellations, we registered the atlases into native space and subsequently performed all analyses in native space.

#### Additional data

All other data were analyzed using Python (Python Software Foundation, version 3.8). For the analysis of the visual attention task, we employed a repeated measures ANOVA from the *statsmodels* module to evaluate effects of condition on attention. The predictor variable was reaction time in seconds, focusing on changes in processing speed, as suggested by previous literature (Graveling et al., 2013a). For analyses of the memory data, we applied the non-parametric Friedman’s test (Friedman, 1937) to investigate whether the condition had a significant impact on memory performance. Memory scores were recorded as the number of words remembered. The learning score was calculated as the average performance on the two immediate recall tasks. For the delayed recall tasks (20 minutes and 24 hours after learning), we used a generalized linear model (GLM; Nelder & Wedderburn, 1972) and included the learning score as a predictor for task performance.

Similarly, blood parameters were analyzed using a repeated measures ANOVA or Friedman’s test, depending on adherence of the data to statistical assumptions. Post-hoc analyses were conducted using paired t-tests or the Wilcoxon signed-rank test (Wilcoxon, 1945), based on the primary statistical method. Blood values during MR acquisition were averaged from time points 48 and 72 minutes (Fig. S1-S6). Missing data (e.g. due to issues in the analysis of the samples) were handled by replacing any missing values for a given subject and condition with the median of the corresponding blood parameter from the other subjects in that condition.

## Data availability

All behavioral data, blood data, and subject-wise, region-averaged CMR_O2_ and CBF data are available as part of our github repository https://github.com/NeuroenergeticsLab/brain_body_metabolism. These data allow the direct replication of the reported mixed model results. To protect participant privacy, the raw imaging and blood data used in the mqBOLD approach are available from the authors upon request.

## Code availability

The code to reproduce all figures and statistical analyses can be found in our GitHub repository: https://github.com/NeuroenergeticsLab/brain_body_metabolism.

## Acknowledgements

We would like to thank Prof. Dr. Christine Preibisch for the provision of MRI protocols as well as template code for the acquisition and processing of mqBOLD MRI data. We are also grateful to Prof. Dr. Claus Zimmer and Prof. Dr. Marcus Makowski for their support in making MR scanning time from the clinic available for our research. Finally, we sincerely thank Dr. Kourosh Zolghadr for his kind support in blood gas analyses.

## Author contributions

AB performed design conceptualization and implementation, investigation, formal analysis, visualization and writing. SJH, JK and FAH performed glucose clamping and provided medical supervision. SM and LS were managed participant handling and assisted with glucose clamping and MR scanning. RI contributed to project conceptualization with glucose clamping and provided medical supervision. VR provided supervision, project conceptualization (neuroimaging and behavioral data), funding and writing.

