## Supplementary material for "The selfish yet forgetful brain: Stable cerebral oxygen metabolism during hypoglycemia but impaired memory consolidation"

by Bose A et al.

**Table S1**

Additional subject characteristics

| **Variable** | **Mean ±sd** |
|---|---|
| Age | 24.03 ±2.1 years |
| Weight | 77.34 ±7.15 kg |
| BMI | 22.93 ±2.46 kg/$m^2$ |
| Body fat | 18.22 ±4.61% |
| Fasting blood glucose | 87.33 ±4.80 mg/dL |
| Fasting tissue glucose | 86.55 ±7.53 mg/dL |
| Hypoglycemic blood glucose | 49.66 ±8.92 mg/dL |
| Artificial euglycemic blood glucose | 91.75 ±7.95 mg/dL |
| Natural euglycemic blood glucose | 88.51 ±4.44 mg/dL |

**Table S2**

Imaging parameters, mean ±SD across subjects and conditions within GM

| $CMR_{O2}$ [µmol/100g/min] | CBF [ml/100g/min] | OEF [ratio] | CBV [%] |
|---|---|---|---|
| 141.68 (19.66) | 49.84 (6.82) | 0.38 (0.03) | 4.79 (0.18) |

*Note.* These values are based on the parameter thresholds mentioned in the methods section.

**Table S3**

Statistical results for blood parameter differences between conditions per timepoint

| Parameter | Timepoint | Statistical test | Test statistic, p-value |
|---|---|---|---|
| Cortisol | Baseline (t=0) | RM ANOVA | F=0.16, p=0.85 |
| | 24 min (t=24) | Friedman’s test | $\chi$2F=1.88, p=0.39 |
| | MRI (t=48, t=72) | RM ANOVA | F=0.65, p=0.53 |
| | Cognitive testing (t=96) | Friedman’s test | $\chi$2F=1.53, p=0.47 |
| **C-peptide** | Baseline (t=0) | Friedman’s test | $\chi$2F=3.57, p=0.17 |
| | 24 min (t=24) | RM ANOVA | F=2.45, p=0.10 |
| | **MRI (t=48, t=72)** | Friedman’s test | $\chi$2F=14.39, p<0.001 |
| | | **Wilcoxon SR (glc)** | **W=9.0, p<0.01** |
| | | Wilcoxon SR (ins) | W=46.5, p=0.27 |
| | **Cognitive testing** (t=96) | Friedman’s test | $\chi$2F=18.37, p<0.001 |
| | | Wilcoxon SR (glc) | W=36.0, p=0.30 |
| | | **Wilcoxon SR (ins)** | **W=11.5, p<0.001** |
| **Epinephrine** | Baseline (t=0) | Friedman’s test | $\chi$2F=2.23, p=0.33 |
| | 24 min (t=24) | Friedman’s test | $\chi$2F=0.23, p=0.89 |
| | **MRI (t=48, t=72)** | Friedman’s test | $\chi$2F=19.78, p<0.001 |
| | | **Wilcoxon SR (glc)** | **W=3.0, p<0.001** |
| | | Wilcoxon SR (ins) | W=46.0, p=0.43 |
| | Cognitive testing (t=96) | Friedman’s test | $\chi$2F=4.79, p=0.09 |
| IGF-1 | Baseline (t=0) | Friedman’s test | $\chi$2F=0.12, p=0.94 |
| | 24 min (t=24) | Friedman’s test | $\chi$2F=1.53, p=0.47 |
| | MRI (t=48, t=72) | RM ANOVA | F=0.01, p=0.99 |
| | Cognitive testing (t=96) | Friedman’s test | $\chi$2F=0.12, p=0.94 |
| **Insulin** | Baseline (t=0) | Friedman’s test | $\chi$2F=7.73, p=0.02 |
| | | Wilcoxon SR (glc) | W=38.0, p=0.07 |
| | | Wilcoxon SR (ins) | W=48.0, p=0.19 |
| | **24 min (t=24)** | Friedman’s test | $\chi$2F=25.53, p<0.001 |
| | | Wilcoxon SR (glc) | W=72.0, p=0.85 |
| | | **Wilcoxon SR (ins)** | **W=0, p<0.001** |
| | **MRI (t=48, t=72)** | Friedman’s test | $\chi$2F=25.53, p<0.001 |
| | | Wilcoxon SR (glc) | W=61.0, p=0.49 |
| | | **Wilcoxon SR (ins)** | **W=0, p<-0.001** |
| | **Cognitive testing** (t=96) | Friedman’s test | $\chi$2F=25.53, p<0.001 |
| | | **Wilcoxon SR (glc)** | **W=23.0, p<0.01** |

| | | | |
|---|---|---|---|
| | | **Wilcoxon SR (ins)** | **W=0, p<0.001** |
| Norepinephrine | Baseline (t=0) | RM ANOVA | F=0.13, p=0.88 |
| | 24 min (t=24) | RM ANOVA | F=2.19, p=0.13 |
| | MRI (t=48, t=72) | RM ANOVA | F=2.28, p=0.12 |
| | Cognitive testing (t=96) | Friedman's test | $\chi$2F=3.29, p=0.19 |

*Note.* Timepoints are provided in minutes since clamp begin at t=0. The timepoint t=MR represents blood samples drawn during MRI acquisition, i.e. die average of plasma concentrations measured at t=48 and t=72. RM ANOVA ≙ repeated measures ANOVA; Wilcoxon SR (glc/ins) ≙ Wilcoxon signed rank test for the glucose or insulin contrast.

§*Please note:* Fig. S1-S6 depict blood parameters across the experiment per condition. Measurements at x=0 reflect baseline levels, acquired before clamping was initiated. Timepoints x=48 and x=72 are used for assessment of blood value differences during the MR scan. After x=72, clamping was stopped. At x=96, previous euglycemia had been restored. At this point, cognitive tests were administered. Contrasts tested were *hypo* vs. $eu_{art}$ (glucose contrast) and $eu_{nat}$ vs. $eu_{art}$ (insulin contrast). Significant levels: <0.001: ***, <0.01: **, <0.05: *. Shown are mean values per condition, together with 95% confidence intervals.

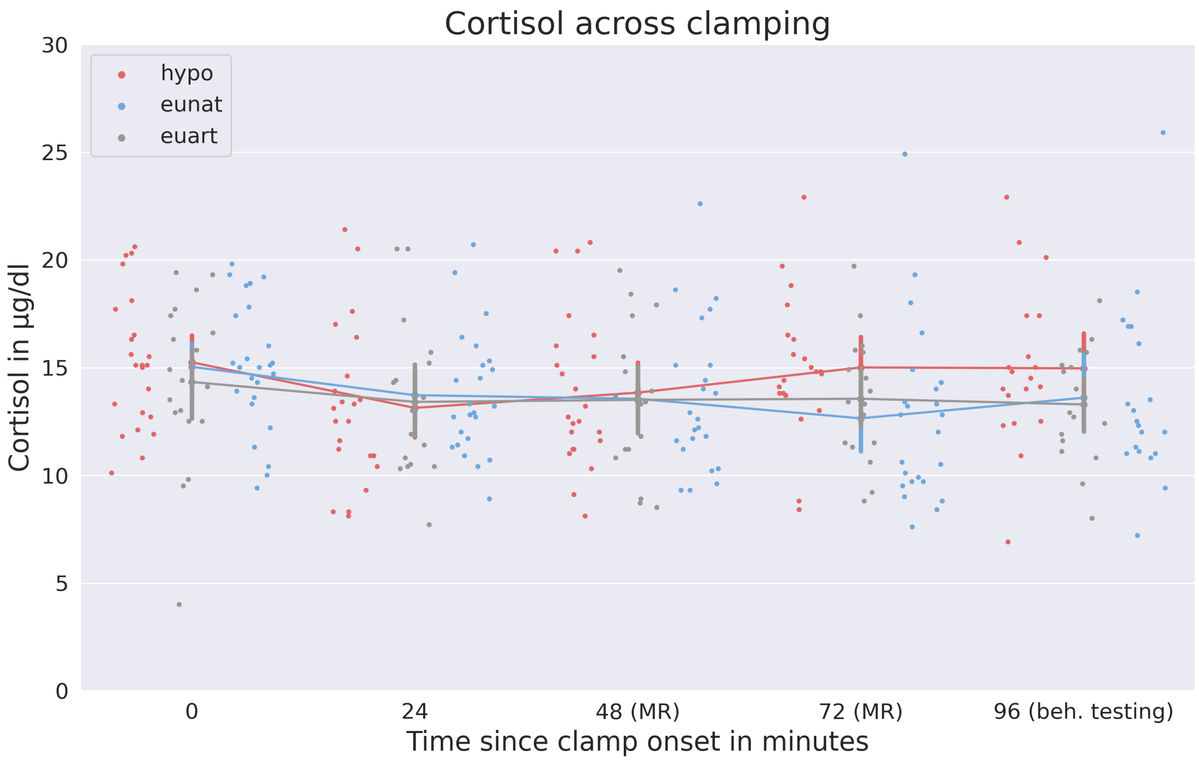


**Fig. S1 | Cortisol levels across the experiment per condition.**§ Cortisol levels did not differ between conditions at any timepoint.

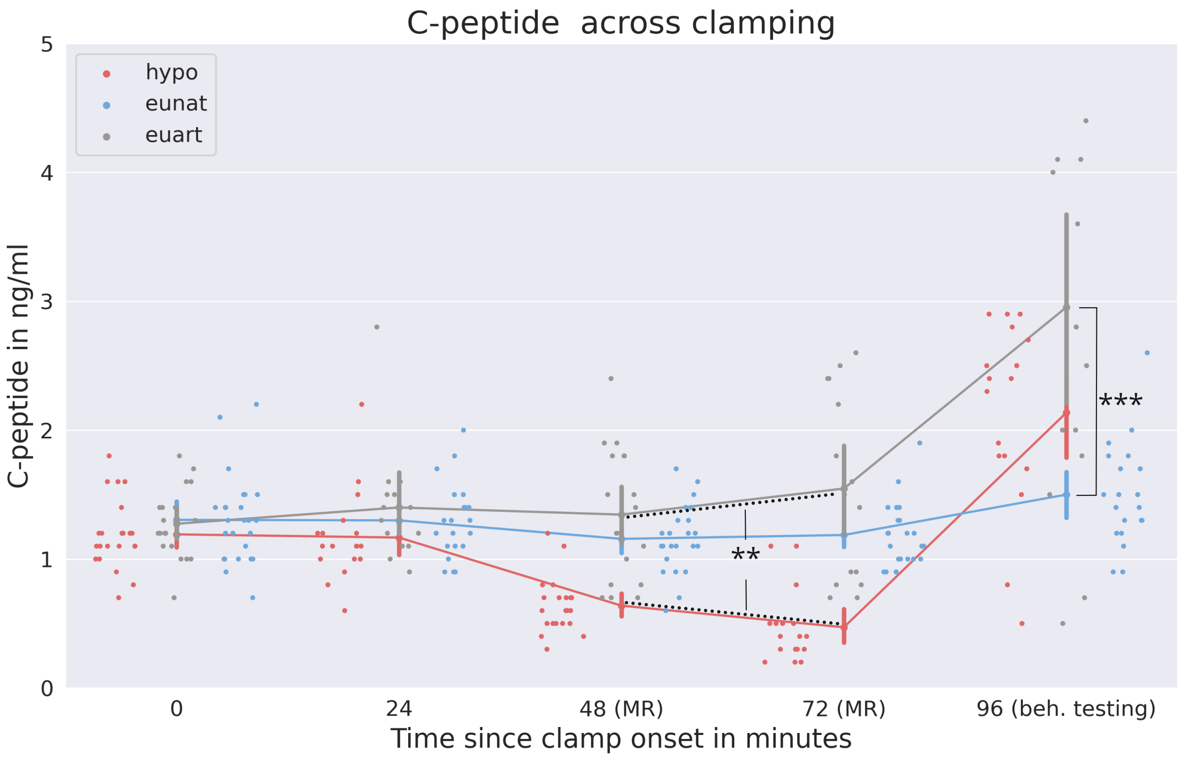


**Fig. S2 | C-peptide levels across the experiment per condition.**§ During the MR scan, c-peptide was significantly lower in *hypo* vs. $eu_{art}$ (glucose contrast). After clamping (x=96), c-peptide was significantly higher in $eu_{art}$ compared to $eu_{nat}$ (insulin contrast).

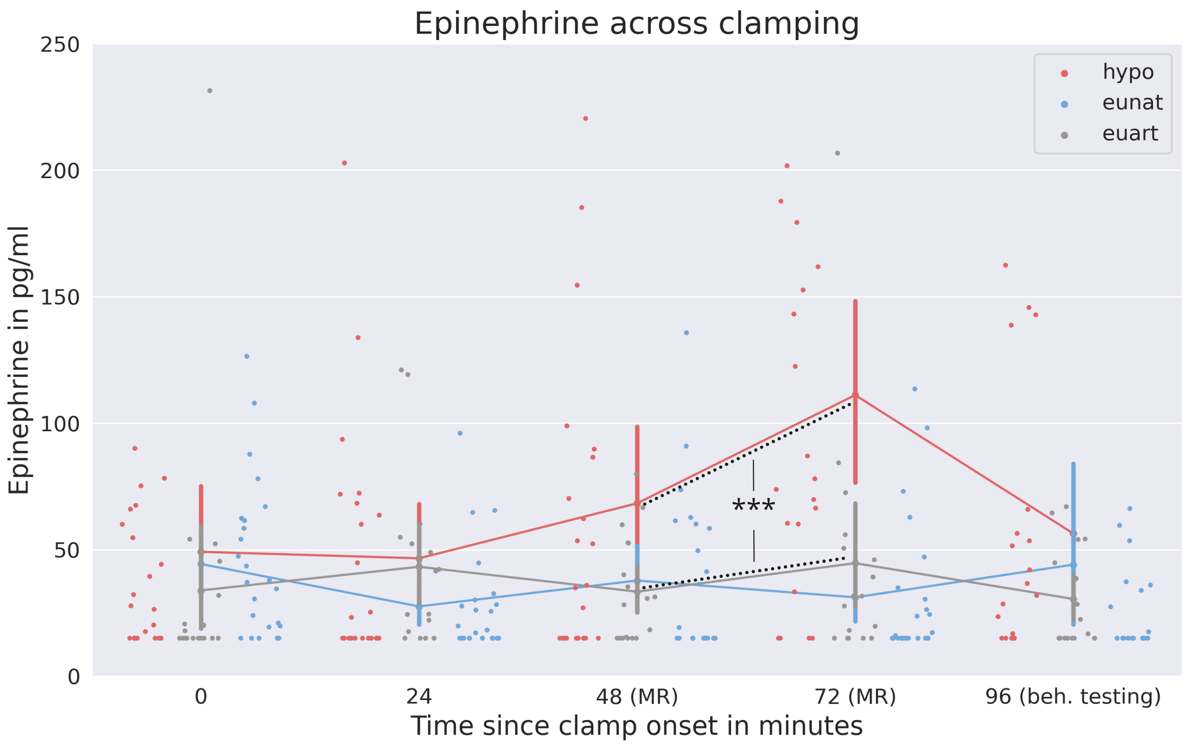


**Fig. S3 | Epinephrine levels across the experiment per condition.**[§] During the MR scan, epinephrine was significantly higher in *hypo* vs. $eu_{art}$ (glucose contrast).

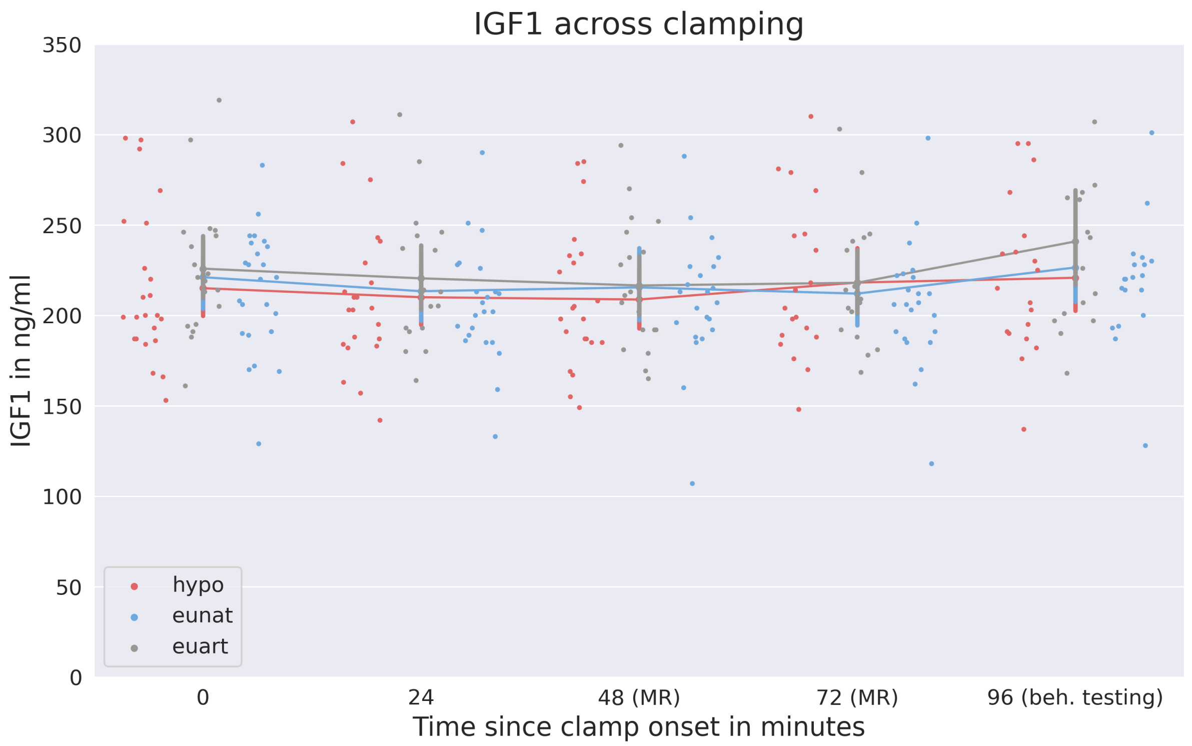


**Fig. S4 | IGF-1 levels across the experiment per condition.**[§] IGF-1 levels did not differ between conditions at any timepoint.

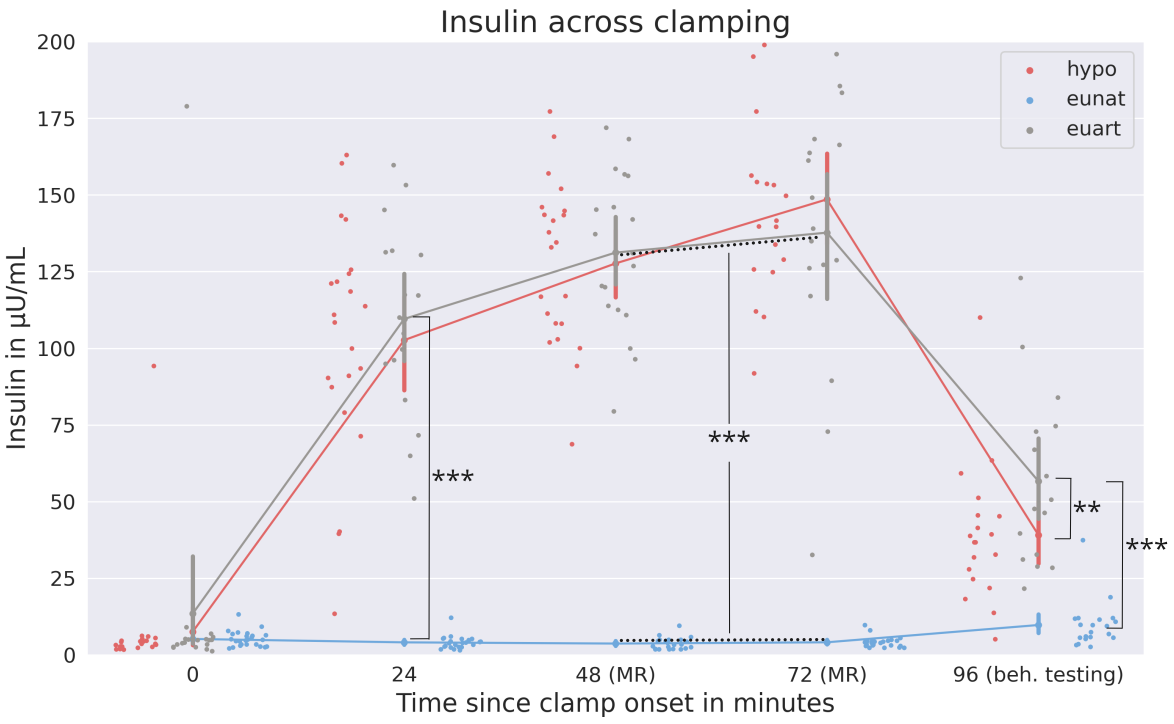


**Fig. S5 | Insulin levels across the experiment per condition.**[§] From t=24 onwards, insulin was significantly higher in $eu_{art}$ than $eu_{nat}$ (insulin contrast). At t=96, insulin was higher in $eu_{art}$ than *hypo* (glucose contrast).

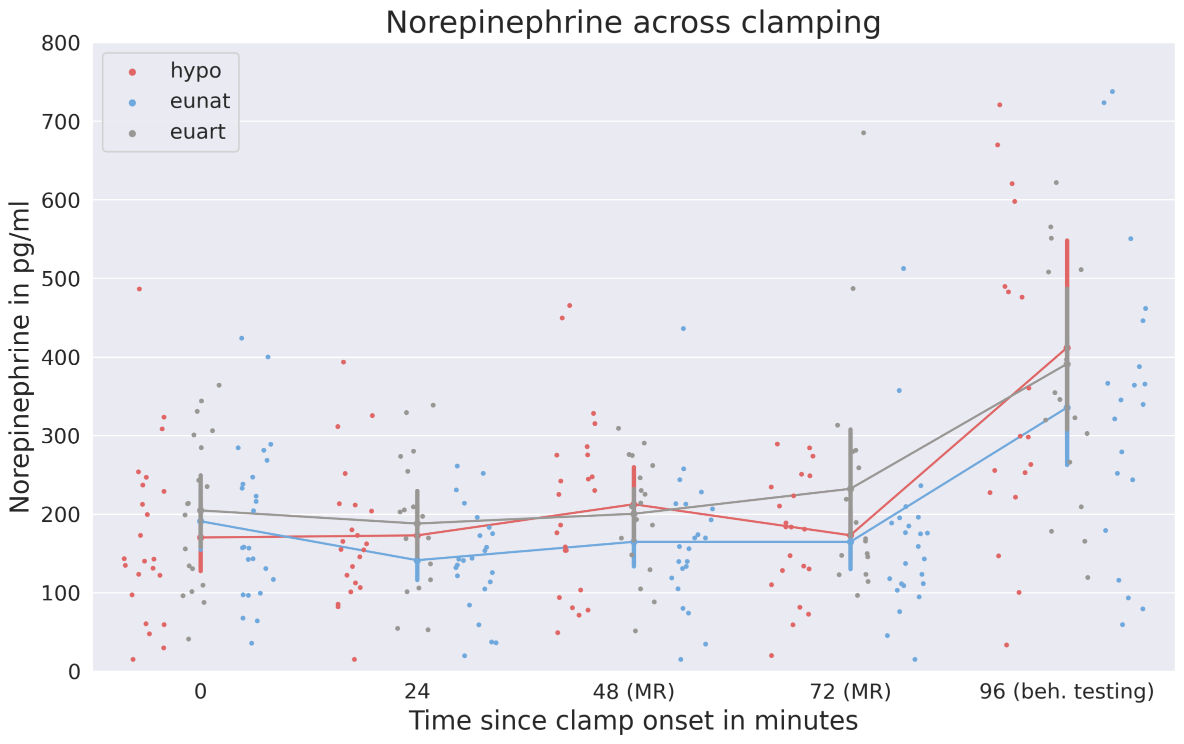


**Fig. S6 | Norepinephrine levels across the experiment per condition.**[§] Norepinephrine levels did not differ between conditions at any timepoint.

**Table S4**

Results of the linear model predicting $CMR_{O2}$ in the main group

| Predictor | Estimate µmol/100g/min | t | 95% CI | p |
|---|---|---|---|---|
| (Intercept) | 130.58 | 30.48 | 121.99 – 139.13 | <0.001*** |
| **Condition (ref: euart)** | | | | |
| Condition[eunat] | -3.34 | -0.93 | -10.49 – 3.79 | 0.35 |
| Condition[hypo] | -0.73 | -0.2 | -8.08 – 6.62 | 0.84 |
| **Network (ref: visual)** | | | | |
| network[Cont] | 19.76 | 11.06 | 16.26 – 23.26 | <0.001*** |
| network[Default] | 15.23 | 9.74 | 12.16 – 18.29 | <0.001*** |
| network[DorsAttn] | 1.58 | 0.86 | -2.03 – 5.18 | 0.39 |
| network[SalVentAttn] | -9.42 | -5.15 | -13.01 – -5.83 | <0.001*** |
| Network[SomMot] | -6.19 | -3.83 | -9.35 – -3.02 | <0.001*** |
| **Interaction condition:network** | | | | |
| Condition[eunat]:network[Cont] | 0.35 | 0.15 | -4.30 – 4.99 | 0.88 |
| Condition[hypo]:network[Cont] | -0.43 | -0.18 | -5.16 – 4.30 | 0.86 |
| Condition[eunat]:network[Default] | 1.39 | 0.67 | -2.68 – 5.45 | 0.50 |
| Condition[hypo]:network[Default] | 2.51 | 1.19 | -1.63 – 6.65 | 0.23 |
| Condition[eunat]:network[DorsAttn] | 3.14 | 1.29 | -1.65 – 7.93 | 0.20 |
| Condition[hypo]:network[DorsAttn] | -1.76 | -0.71 | -6.63 – 3.12 | 0.48 |
| Condition[eunat]:network[SalVentAttn] | 0.94 | 0.39 | -3.82 – 5.70 | 0.70 |
| Condition[hypo]:network[SalVentAttn] | 0.23 | 0.09 | -4.62 – 5.08 | 0.93 |
| Condition[eunat]:network[SomMot] | 1.86 | 0.87 | -2.34 – 6.07 | 0.39 |
| Condition[hypo]:network[SomMot] | 0.20 | 0.09 | -4.08 – 4.49 | 0.93 |

*Note.* Result parameters for the main group of the following model: $CMR_{O2}$ ~ condition*network + (1|subject/condition). Marginal $R^2$ / conditional $R^2$ = 5.33% / 22.51%. Network abbreviations: Cont ≙ Control; Default ≙ Default mode; DorsAttn ≙ Dorsal attention; SalVentAttn ≙ Salience; SomMot ≙ Somatomotor. Significant codes: <0.001: ***; <0.01: **, <0.05: *. Analyses were performed in native space.

**Table S5**

Results of the linear model predicting $CMR_{O2}$ in the subgroup

| Predictor | Estimate µmol/100g/min | t | 95% CI | p |
|---|---|---|---|---|
| (Intercept) | 130.29 | 28.99 | 121.17 - 139.35 | <0.001*** |
| **Condition (ref: euart)** | | | | |
| Condition[hypo] | -0.67 | -0.12 | -11.60 - 10.25 | 0.90 |
| **Network (ref: visual)** | | | | |
| network[Cont] | 19.76 | 11.02 | 16.24 - 23.27 | <0.001*** |
| network[Default] | 15.23 | 9.71 | 12.15 - 18.30 | <0.001*** |
| network[DorsAttn] | 1.58 | 0.85 | -2.05 - 5.20 | 0.39 |
| network[SalVentAttn] | -9.42 | -5.13 | -13.02 - -5.82 | <0.001*** |
| network[SomMot] | -6.19 | -3.81 | -9.37 - -3.01 | <0.001*** |
| **Interaction condition:network** | | | | |
| Condition[hypo]:network[Cont] | 2.36 | 0.69 | -4.39 - 9.12 | 0.49 |
| Condition[hypo]:network[Default] | 5.91 | 1.96 | -0.01 - 11.83 | 0.05 |
| Condition[hypo]:network[DorsAttn] | 0.17 | 0.05 | -6.81 - 7.15 | 0.96 |
| Condition[hypo]:network[SalVentAttn] | 0.43 | 0.12 | -6.51 - 7.37 | 0.90 |
| Condition[hypo]:network[SomMot] | 2.85 | 0.91 | -3.28 - 8.97 | 0.36 |

*Note.* Result parameters for the sub group of the following model: $CMR_{O2}$ ~ condition*network + (1|subject/condition). Marginal $R^2$ / conditional $R^2$ = 5.49% / 22.26%. Network abbreviations: Cont ≙ Control; Default ≙ Default mode; DorsAttn ≙ Dorsal attention; SalVentAttn ≙ Salience; SomMot ≙ Somatomotor. Significant codes: <0.001: ***; <0.01: **, <0.05: *. Analyses were performed in native space.

**Table S6**

Results of the linear model predicting CBF in the main group

| Predictor | Estimate µmol/100g/min | t | 95% CI | p |
|---|---|---|---|---|
| (Intercept) | 43.56 | 26.38 | 40.27 - 46.87 | <0.001*** |
| **Condition (ref: euart)** | | | | |
| Condition[eunat] | -1.23 | -0.81 | -4.23 - 1.80 | 0.42 |
| Condition[hypo] | 0.33 | 0.22 | -2.74 - 3.56 | 0.83 |
| **Network (ref: visual)** | | | | |
| network[Cont] | 14.50 | 34.12 | 13.67 - 15.34 | <0.001*** |
| network[Default] | 10.11 | 27.13 | 9.38 - 10.84 | <0.001*** |
| network[DorsAttn] | 8.31 | 18.89 | 7.45 - 9.17 | <0.001*** |
| network[SalVentAttn] | 8.97 | 20.52 | 8.11 - 9.83 | <0.001*** |
| network[SomMot] | 8.25 | 21.37 | 7.49 - 9.01 | <0.001*** |
| **Interaction condition:network** | | | | |
| Condition[eunat]:network[Cont] | 0.75 | 1.39 | -0.36 - 1.85 | 0.18 |
| Condition[hypo]:network[Cont] | 2.29 | 3.99 | 1.16 - 3.41 | <0.001*** |
| Condition[eunat]:network[Default] | 0.74 | 1.51 | -0.23 - 1.71 | 0.13 |
| Condition[hypo]:network[Default] | 1.94 | 3.84 | 0.95 - 2.92 | <0.001*** |
| Condition[eunat]:network[DorsAttn] | 0.94 | 1.61 | -0.21 - 2.08 | 0.11 |
| Condition[hypo]:network[DorsAttn] | 0.98 | 1.64 | -0.19 - 2.14 | 0.10 |
| Condition[eunat]:network[SalVentAttn] | 0.34 | 0.59 | 0.79 - 1.48 | 0.55 |
| Condition[hypo]:network[SalVentAttn] | 1.20 | 2.02 | 0.04 - 2.35 | 0.04* |
| Condition[eunat]:network[SomMot] | 0.45 | 0.88 | -0.56 - 1.45 | 0.38 |
| Condition[hypo]:network[SomMot] | 0.99 | 1.91 | -0.28 - 2.02 | 0.06 |

*Note.* Result parameters for the main group of the following model: CBF ~ condition*network + (1|subject/condition). Marginal $R^2$ / conditional $R^2$ = 12.24% / 44.70%. Network abbreviations: Cont ≙ Control; Default ≙ Default mode; DorsAttn ≙ Dorsal attention; SalVentAttn ≙ Salience; SomMot ≙ Somatomotor. Significant codes: <0.001: ***; <0.01: **, <0.05: *. Analyses were performed in native space.

**Table S7**

Results of the linear model predicting CBF in the subgroup

| Predictor | Estimate µmol/100g/min | t | 95% CI | p |
|---|---|---|---|---|
| (Intercept) | 43.08 | 26.74 | 39.83 - 46.33 | <0.001*** |
| **Condition (ref: euart)** | | | | |
| Condition[hypo] | 3.12 | 1.44 | -1.28 - 7.48 | 0.16 |
| **Network (ref: visual)** | | | | |
| network[Cont] | 14.50 | 33.50 | 13.65 - 15.35 | <0.001*** |
| network[Default] | 10.11 | 26.64 | 9.37 - 10.85 | <0.001*** |
| network[DorsAttn] | 8.31 | 18.55 | 7.43 - 9.19 | <0.001*** |
| network[SalVentAttn] | 8.97 | 20.15 | 8.10 - 9.84 | <0.001*** |
| network[SomMot] | 8.25 | 20.98 | 7.48 - 9.02 | <0.001*** |
| **Interaction: condition:network** | | | | |
| Condition[hypo]:network[Cont] | 4.79 | 5.74 | 3.15 - 6.42 | <0.001*** |
| Condition[hypo]:network[Default] | 3.89 | 5,32 | 2.46 - 5.33 | <0.001*** |
| Condition[hypo]:network[DorsAttn] | 2.03 | 2.35 | 0.34 - 3.72 | 0.02* |
| Condition[hypo]:network[SalVentAttn] | 1.93 | 2.25 | 0.25 - 3.61 | 0.02* |
| Condition[hypo]:network[SomMot] | 1.63 | 2.15 | 0.15 - 3.12 | 0.03* |

*Note.* Result parameters for the subgroup of the following model: CBF ~ condition*network + (1|subject/condition). Marginal $R^2$ / conditional $R^2$ = 14.05% / 42.27%. Network abbreviations: Cont ≙ Control; Default ≙ Default mode; DorsAttn ≙ Dorsal attention; SalVentAttn ≙ Salience; SomMot ≙ Somatomotor. Significant codes: <0.001: ***; <0.01: **, <0.05: *. Analyses were performed in native space.

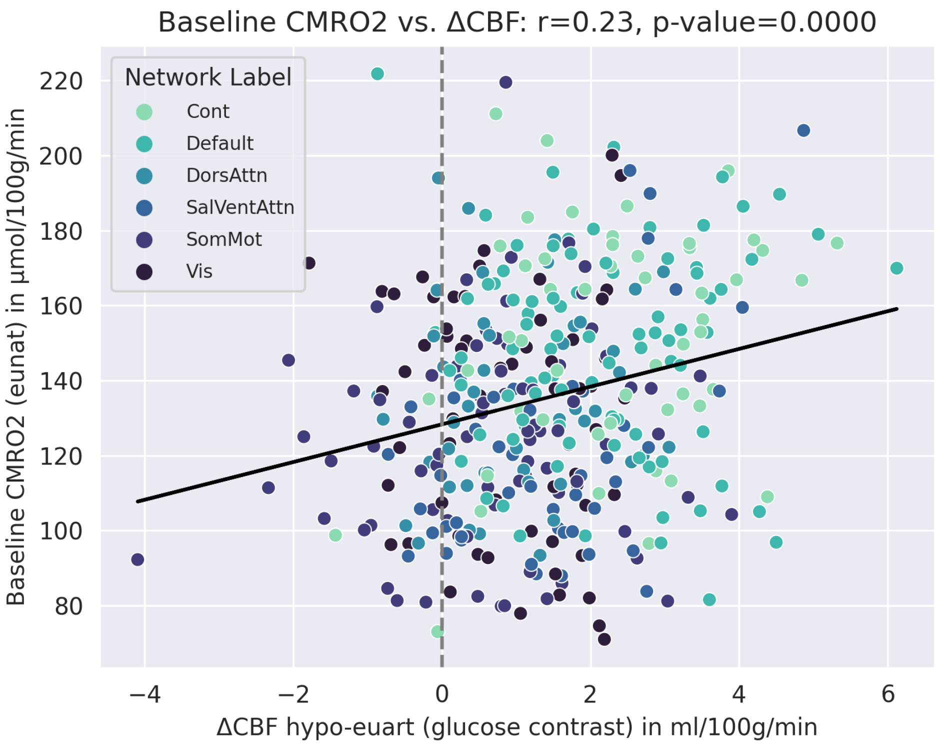


**Fig. S7 |Correlations between baseline metabolism (CMRO2 in eunat) and CBF increase in the glucose contrast (hypo-euart) per ROI, color-coded by network.** The black line represents the line of best fit. CBF increase in hypo correlates positively with baseline metabolism (r=0.23, p<0.001). Network abbreviations: Cont ≙ Control; Default ≙ Default mode; DorsAttn ≙ Dorsal attention; SalVentAttn ≙ Salience; SomMot ≙Somatomotor. Analyses were performed in native space.
